# PBS3 is the missing link in plant-specific isochorismate-derived salicylic acid biosynthesis

**DOI:** 10.1101/600692

**Authors:** Dmitrij Rekhter, Daniel Lüdke, Yuli Ding, Kirstin Feussner, Krzysztof Zienkiewicz, Volker Lipka, Marcel Wiermer, Yuelin Zhang, Ivo Feussner

**Affiliations:** University of Goettingen, Albrecht-von-Haller-Institute for Plant Sciences, Department of Plant Biochemistry, Justus-von-Liebig Weg11, D-37077 Goettingen, Germany.; University of Goettingen, Albrecht-von-Haller-Institute for Plant Sciences, RG Molecular Biology of Plant-Microbe Interactions, Julia-Lermontowa-Weg 3, D-37077 Goettingen, Germany.; University of British Columbia, Department of Botany, Vancouver, BC V6T 1Z4, Canada.; University of Goettingen, Goettingen Center for Molecular Biosciences (GZMB), Service Unit for Metabolomics and Lipidomics, Justus-von-Liebig Weg11, D-37077 Goettingen, Germany.; University of Goettingen, Albrecht-von-Haller-Institute for Plant Sciences, Department of Plant Cell Biology, Julia-Lermontowa-Weg 3, D-37077 Goettingen, Germany.; University of Goettingen, Central Microscopy Facility of the Faculty of Biology & Psychology, Julia-Lermontowa-Weg 3, D-37077 Goettingen, Germany.; University of Goettingen, Goettingen Center for Molecular Biosciences (GZMB), Department of Plant Biochemistry, Justus-von-Liebig Weg11, D-37077 Goettingen, Germany.

## Abstract

The phytohormone salicylic acid (SA) is a central regulator of plant immunity. Despite such functional importance, our knowledge of its biosynthesis is incomplete. Previous work showed that SA is synthesized from chorismic acid in plastids. The bulk of pathogen-induced SA derives from isochorismate generated by the catalytic activity of ISOCHORISMATE SYNTHASE1 (ICS1). How and in which cellular compartment isochorismate is converted to SA is unknown. Here we show that the pathway downstream of isochorismate requires only two additional proteins: the plastidial isochorismate exporter ENHANCED DISEASE SUSCEPTIBILITY5 (EDS5) and the cytosolic amido-transferase AvrPphB SUSCEPTIBLE3 (PBS3). PBS3 catalyzes the conjugation of glutamate to isochorismate. The reaction product isochorismate-9-glutamate spontaneously decomposes into enolpyruvyl-*N*-glutamate and SA. This previously unknown reaction mechanism appears to be conserved throughout the plant kingdom.

**One Sentence Summary:** Salicylic acid is synthesized via isochorismate-9-glutamate by PBS3.

## Main Text

The phenolic plant hormone salicylic acid (SA) is a central signaling molecule required for adaptive responses to biotic and abiotic stress conditions (*1*). In marked contrast to the functional relevance of SA, a detailed understanding of its plant-specific biosynthesis pathway(s) is still incomplete (*2*). Generally, plants can produce SA via two distinct metabolic processes, both starting in plastids with the branch-point metabolite chorismic acid. In the model plant *Arabidopsis thaliana*, only a minor proportion of defense-related SA is produced by the phenylpropanoid/PAL-pathway, whereas the majority (c. 90%) is derived from isochorismate that is generated by the plastid-localized ISOCHORISMATE SYNTHASE1 (ICS1) (Fig. 1A) (*3–5*). In order to execute its signaling functions, either SA or one of its precursors needs to be transported out of the plastids (*6*). The MATE family transporter ENHANCED DISEASE SUSCEPTIBILITY5 (EDS5) localized in the chloroplast envelope is required in this process (*7*). Except for ICS1 and EDS5, molecular components required for completion of the ICS1-dependent biosynthetic route of SA remain enigmatic. SA-producing bacteria employ an isochorismate pyruvate lyase (IPL) to directly convert isochorismate to SA (*8*). However, *IPL* homologues have not been found in any plant genome so far and how isochorismate is converted to SA is unknown (*9*).

**Fig. 1.**
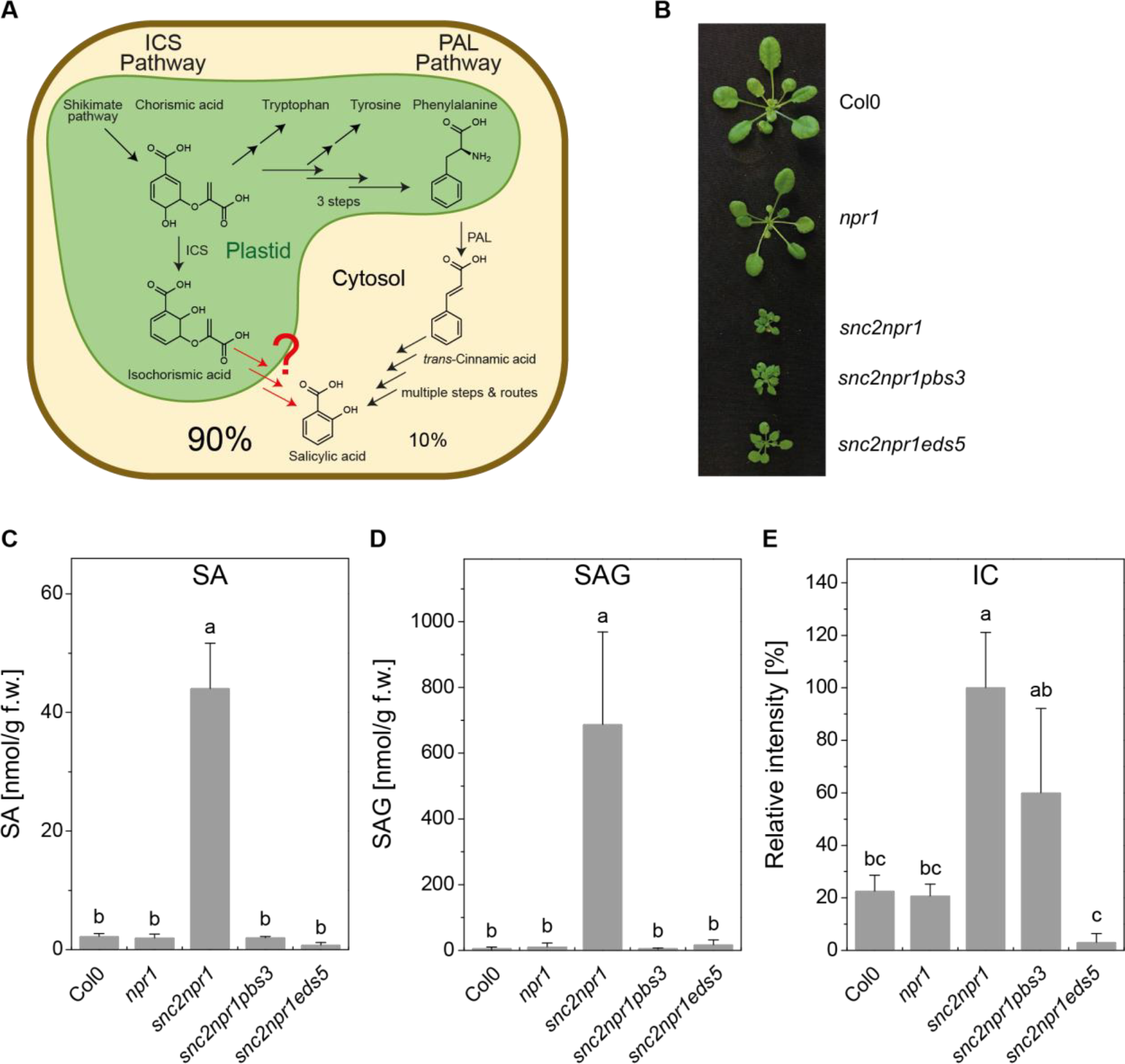
Alternative salicylic acid (SA) biosynthetic routes in plants and analysis of SA-related metabolites in autoimmune mutants. (A) Upon pathogen infection, SA synthesis derives from aromatic amino acid biosynthesis in plastids (Shikimate pathway) and may follow two alternative pathways. PAL pathway: synthesis starts from phenylalanine via cinnamic acid. ICS pathway: starts from chorismate via isochorismate. Analysis of SA formation in *ics1* suggests that at least 90% of the SA is being synthesized from chorismic acid in *Arabidopsis* leaves (*4*). (B-E) Analysis of SA-related metabolites in the indicated plant genotypes. (B) Morphology of four-week-old plants. (C-E) Absolute amounts of SA (B) and SA-glycoside (SAG, D) and relative amounts of isochorismate (IC, E). Absolute and relative amounts were determined by LC-MS analysis of *Arabidopsis* leaf samples. Bars represent the mean ± STD of three biological replicates. Statistical differences among replicates are labeled with different letters (P < 0.05, one-way ANOVA and post hoc Tukey’s Test; n = 3).

Similar to plants with defects in *ICS1* and *EDS5*, mutants of AvrPphB SUSCEPTIBLE3 (PBS3) also show reduced pathogen-induced SA accumulation (*10, 11*). *PBS3* belongs to the GH3 acyl-adenylase gene family, whose founding member JASMONIC ACID RESPONSE1 (JAR1; AtGH3.11) catalyzes the conjugation of isoleucine to jasmonic acid, yielding the active hormone conjugate jasmonoyl-isoleucine (*12*). Although SA was shown to be a poor substrate of PBS3 (*13*), its transcriptional co-regulation with *ICS1* and *EDS5* (Fig. S1) supports a coordinated contribution to SA or SA-precursor biosynthesis, modification or transport. A central component of canonical SA signaling is *NONEXPRESSOR OF PATHOGENESIS-RELATED GENES 1* (*NPR1*) which functions as an SA-dependent regulator of defense gene expression (*14*). Consequently, *npr1* mutant plants show compromised defense responses to a wide range of pathogens (*15*). A forward genetic suppressor screen in the *npr1*-1 mutant background previously identified a gain-of-function mutation in the receptor-like protein SNC2. This *snc2*-1D (for *suppressor of npr1*-1*, constitutive 2*) mutant shows an autoimmune phenotype accompanied by 20-fold higher constitutive amount of SA and stunted growth (*16*) (Fig. 1B and C). We used this mutant as a tool to investigate the contribution of PBS3 and EDS5 to SA accumulation in corresponding triple mutant combinations in *snc2-*1*D npr1-*1 background (Figure 1 C to E).

Both triple mutants *snc2-*1D *npr1-*1 *pbs3-*1 and *snc2-*1*D npr1-*1 *eds5-*3 retained the dwarf growth phenotype (Fig. 1B), which can be explained by constitutive activation of not only SA-dependent, but also SA-independent resistance pathways in the *snc2-*1D *npr1*-1 mutant (*16*). However, metabolite analyses revealed that the elevated accumulation of both SA (20-fold) and its glycosylated storage form SA-glycoside (SAG, 120-fold) are reduced back to wild-type levels in both triple mutants (Fig. 1C, D). In marked contrast, the ICS1 reaction product isochorismate accumulated 3-fold higher over wild-type in the *snc2-*1D *npr1-*1 *pbs3-* 1 triple mutant (Fig. 1E). Based on this observation, we hypothesized that isochorismate is the substrate of PBS3. Isochorismate did not accumulate in the *snc2-*1D *npr1-*1 *eds5-*3 mutant, most likely because ICS1 operates near equilibrium and isochorismate cannot be exported from the plastid by EDS5 (*17*).

To test our hypothesis, we purified recombinant PBS3 and ICS1 to homogeneity (Figs. S2 and S3). As isochorismate is commercially not available, we used ICS1 to produce isochorismate from chorismate (Fig. S4). This isochorismate preparation included residual amounts of chorismate and was incubated with PBS3 (Fig. 2D). The reaction monitored by liquid chromatography coupled to accurate mass high resolution mass spectrometry yielded four signals for the extracted ion chromatogram of *m/z* 354.083 (Fig. 2A black line). Two minor signals represented chorismate-conjugates (Fig. 2A black line, a and d, 2B a and d, S5A - D). The signal at 2.85 min was identified as isochorismate-9-glutamate by accurate mass fragmentation analyses (Fig. 2A black line c, 2B c, S6B - D). The fourth signal belonged to isochorismate-7-glutamate (Fig. 2A black line b, 2B b, S6E and F). Based on the signal intensities, we concluded that isochorismate-9-glutamate was the preferred product of PBS3. The incubation of PBS3 with chorismate confirmed the structure of two minor products (Fig. 2A red line a and d) (*13*). It was previously described that isochorismate itself decomposes in aqueous solution into SA and enolpyruvate (*18*). We confirmed this observation by detecting small amounts of SA in the absence of either glutamate, ATP or PBS3 in the assay after one hour of incubation (Fig. 2C, colored lines). However, in the presence of PBS3 we detected four times more SA beside the glutamate conjugates (Fig. 2C, black line). This difference is most likely higher *in planta*, because we detected non-enzymatic decomposition of isochorismate into SA before adding PBS3 to the reaction mixture (Fig. 2C and F). As no lyase activity has been reported for GH3 proteins so far, the formation of isochorismate-9-glutamate most likely accelerated the non-enzymatic decomposition of isochorismate by elimination, leading to the increased accumulation of SA.

**Fig. 2.**
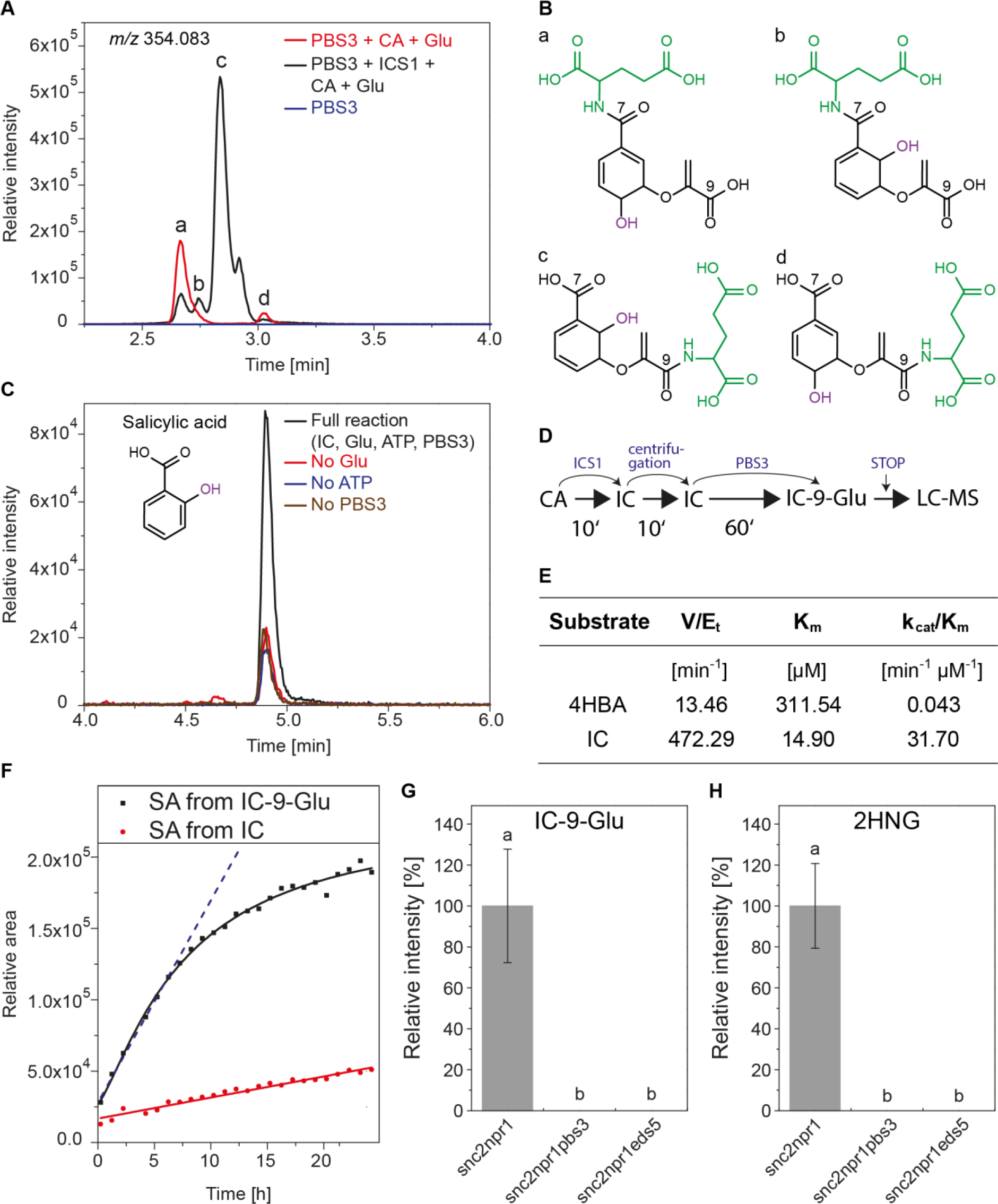
PBS3 catalyzes the formation of isochorismate-9-glutamate, which non-enzymatically decomposes into salicylic acid (SA). (A) LC-MS analysis of products from activity assays with purified recombinant PBS3. The assays were performed by adding chorismate (CA) and glutamate (Glu) (red line), with CA, Glu and ISOCHORISMATE SYNTHASE 1 (ICS1, which converts CA to isochorismate (IC), black line) or without adenosine triphosphate (ATP, blue line). The extracted ion chromatogram for isochorismate-9-glutamate (*m/z* 354.083) is shown. The main signal (c) has an adsorption maximum of 278 nm as previously described for isochorismate (*32*). (B) Chemical structures of conjugates formed by PBS3 with Glu (green) and chorismate (a, d) or isochorismate (b, c). Depending on the substrate, glutamate is preferably conjugated to C7 (for chorismate) or C9 (for isochorismate). Structures were solved with MS/MS (Fig. S5 and S6). (C) Extracted ion chromatogram for SA (*m/z* 137.024) which accumulates in the activity assay with PBS3 in the presence of isochorismate (IC), glutamate (Glu) and ATP (black line), but only in minor amount in the absence of Glu (red line), ATP (blue line) or PBS3 (brown line). The presence of SA in the control assays is due to chemical decomposition of isochorismate in aqueous solution. The identity of SA was confirmed by an authentic standard. (D) Scheme describing experimental time course of the analysis shown in (C). (E) Kinetic parameters of PBS3 with 4-hydroxy benzoate (4HBA) or isochorismate (IC) as substrate. Data were obtained spectrophotometrically in triplicates. (F) Time course for the formation of SA from the non-enzymatic decomposition of isochorismate (IC, red line) and isochorismate-9-glutamate (IC-9-Glu, black line). (G, H) Relative amounts of isochorismate-9-glutamate (IC-9-Glu, F) and 2-hydroxy acryloyl-*N*-glutamic acid (2HNG, H) in *snc2 npr1, snc2 npr1 pbs3* and *snc2 npr1 eds5* plants. Bars represent the mean ± STD of three biological replicates. Statistical differences among replicates are labeled with different letters (P < 0.05, one-way ANOVA and post hoc Tukey’s Test; n = 3).

GH3 proteins are bifunctional enzymes (*12*). They catalyze the adenylation of carboxy groups, which then react with an amino group to form an amide bond. Adenosine monophosphate is released from the intermediate. The kinetic parameters for the reaction of PBS3 with isochorismate and glutamate were determined spectrophotometrically to compare the data with those described previously for 4-hydroxy benzoic acid (Fig. 2E) (*13, 19*). The catalytic efficiency (k_cat_/K_m_) of PBS3 was 737 times higher when isochorismate (31.70 min^-1^ µM^−1^) was used as the carboxy substrate in place of 4-hydroxy benzoic acid (0.043 min^−1^ µM^−1^). Such increase is a result of both a higher substrate affinity and an enhanced turnover rate. This fast reaction rate is of physiological importance, since isochorismate otherwise rearranges chemically eight times faster into isoprephenate than it decomposes into SA (*18*). To additionally confirm that isochorismate is the preferred substrate, it was modeled into the binding pocket of PBS3 (Fig. S7B) using a previously published crystal structure in complex with SA and adenosine monophosphate (PDB ID 4eql, Fig. S7A) (*20*). The superimposition of the ring-structure of SA with that of isochorismate revealed that i) there is space to accommodate isochorismate in the binding pocket (Fig. S7C and D); and ii) the two reaction partners (phosphate group of adenosine monophosphate and the C9-carboxy group of isochorismate) are in close proximity (c. 2.1 Å; Fig. S7B).

Beside SA, 2-hydroxy acryloyl-*N*-glutamic acid (or enolpyruvyl-N-glutamate) was detected (*m/z* 216.051) as the second product of the non-enzymatic decomposition of isochorismate-9-glutamate in the *in vitro* assay mixture (Figs. 2F and S6E and F). This observation was confirmed by detecting isochorismate-9-glutamate and 2-hydroxy acryloyl-*N*-glutamate in *snc2-*1D *npr1-*1, but not in the triple mutant plants, *snc2-*1D *npr1-*1 *pbs3-*1 and *snc2-* 1D *npr1-*1 *eds5-*3 (Figs. 2G and H, S6A). Based on these data, we concluded that isochorismate-9-glutamate is the *in planta* precursor of SA. The rates of the non-enzymatic decomposition of isochorismate-9-glutamate or isochorismate into SA were compared over a 24 h time course. During the first 6 h, the formation of SA from isochorismate-9-glutamate is 10-times higher compared to the formation of SA from isochorismate (Fig. 2F). In order to explain this accelerated decomposition, we employed molecular modeling. The resulting structure of isochorismate-9-glutamate suggested the formation of two additional hydrogen bonds between the two carboxy groups of glutamate and the C2-hydroxy and C7-carboxy group of isochorismate (Fig. S9A) instead of only two as for isochorismate (*18*). Thereby the amide hydrogen of the peptide bond and the oxygen of the ether bridge are brought in closer proximity and this shorter distance is most likely sufficient to enhance the reaction rate of the hydrogen transfer. As a consequence of this protonation, a base-initiated aromatization of the ring system can be expected, followed by an elimination of the E1 type that yields SA and 2-hydroxy acryloyl-*N*-glutamate as final products (Fig. S9B).

The biosynthesis of SA was previously proposed to occur in plastids, as ICS1 was shown to localize to this compartment (*4*) (Fig. 1A). However, sequence analyses utilizing the *in silico* online tools TargetP (*21*) and Predotar (*22*) predicted a cytosolic localization of PBS3, consistent with studies characterizing other GH3 protein family members (*23, 24*). We used *Arabidopsis efr* mutant plant leaves, which allow higher levels of *Agrobacterium*-mediated transformation rates (*25*) for transient co-expression of PBS3 C-terminally tagged with YELLOW FLUORESCENT PROTEIN (PBS3-YFP) and ICS1 fused to CYAN FLUORESCENT PROTEIN (ICS1-CFP). Confocal laser scanning microscopy confirmed the presence of ICS1-CFP in plastids and PBS3-YFP in the cytosol (Fig. 3A). Fluorescence lifetime imaging (FLIM) analyses validated the identity of the corresponding fluorescence signals in the respective cellular compartments (Fig. S10A). Immunoblot analyses demonstrated production of full-length fusion proteins (Fig. S10C). Functionality of the PBS3-YFP fusion protein was independently confirmed by transient expression in the *pbs3*-1 single mutant, which fully restored *Agrobacterium*-induced SA accumulation (Fig. S11). Our biochemical and cell biological data strongly suggest a spatial separation of cytosolic PBS3 and its substrate isochorismate, which is produced by plastid-localized ICS1. We therefore hypothesized that the plastidial envelope-residing EDS5 most likely exports isochorismate and not SA into the cytosol, where it is further metabolized by PBS3 (Fig. 3D). This is supported by the observation that SA does not accumulate in *eds5* mutants (*6*) and that expression of native PBS3 alone or in combination with ICS1 does not restore SA accumulation in *eds5-3* mutant plants (Fig. 3C).

**Fig. 3.**
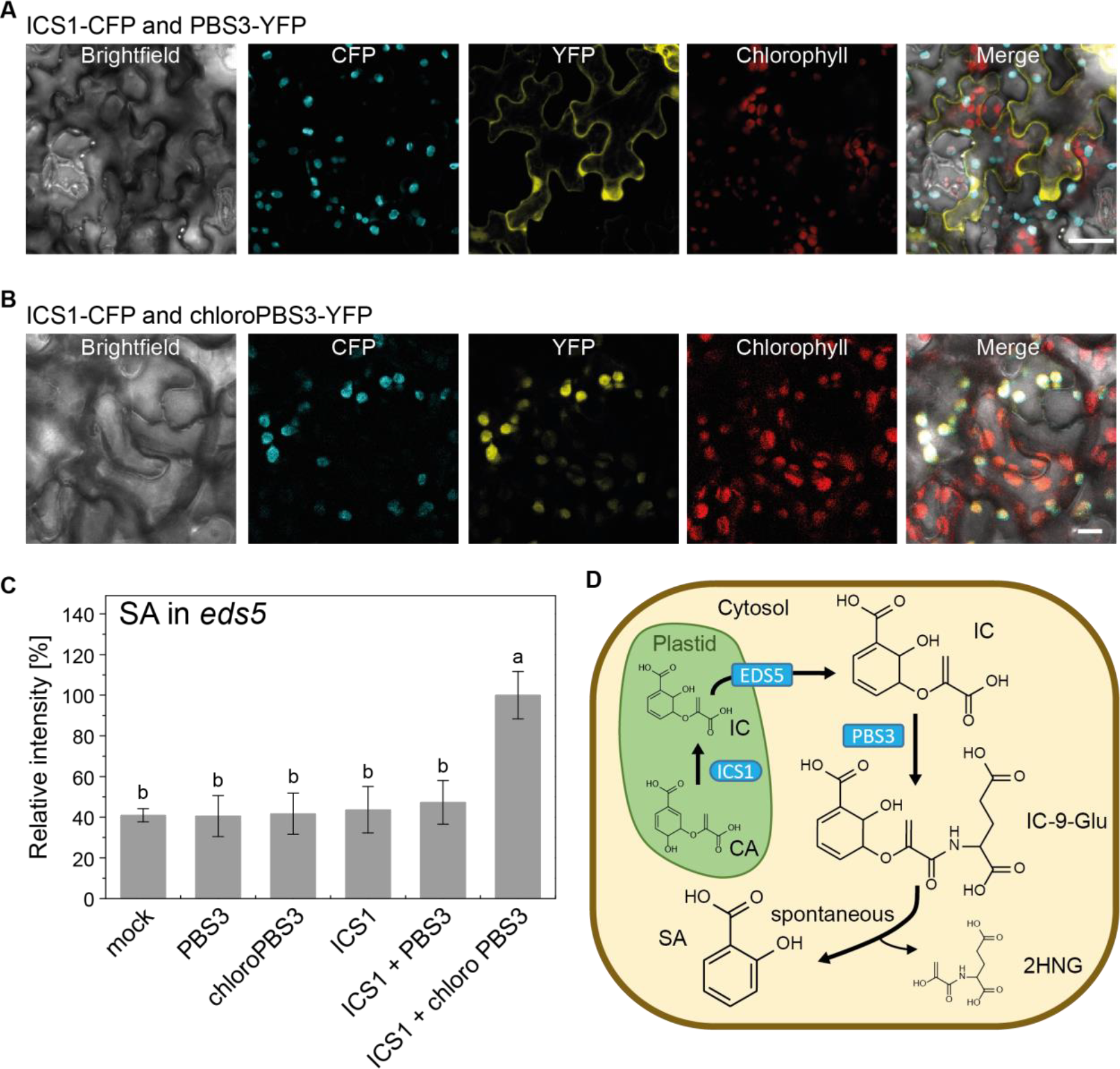
PBS3 and ICS1 are localized in different compartments. Confocal laser scanning microscopy analysis three days after *Agrobacterium*-infiltration for transient co-expression of (A) ICS1-CFP and PBS3-YFP, or (B) ICS1-CFP and chloroPBS3-YFP in *Arabidopsis efr* leaves. Scale bar represents 10 µm. (C) Transient expression of ICS1 and chloroPBS3 in *Arabidopsis eds5-3* leaves restores the *Agrobacterium* induced SA accumulation. 24 h after infiltration, leaves were collected, and metabolites were extracted as described for the metabolite fingerprint analysis. Infiltration medium was used as mock treatment. The SA content was analyzed by LC-MS. Bars represent the mean ± STD of three biological replicates. Statistical differences among replicates are labeled with different letters (P < 0.05, one-way ANOVA and post hoc Tukey’s Test; n = 3). (D) Model summarizing pathogen-induced SA biosynthesis in plants.

We reasoned further that forced mis-targeting of PBS3 into plastids would allow restoration of SA accumulation in the transport-deficient *eds5* mutant. To test this hypothesis, we fused the plastid transit peptide of ICS1 (*4*) to the N-terminus of PBS3-YFP (chloroPBS3-YFP). Transient co-expression of ICS1-CFP and chloroPBS3-YFP in *Arabidopsis efr* mutant leaves confirmed co-localization of the full-length fluorescent fusion proteins in plastids (Fig. 3B; Fig. S10B and C). Most importantly, plastidial co-localization of both ICS1 and PBS3 restored *Agrobacterium*-induced SA synthesis in *Arabidopsis eds5-3* mutant plants (Fig. 3C). Individual expression of ICS1 and PBS3 protein variants or combined expression of ICS1 with cytosolic PBS3 did not. The observation that plastid-targeted PBS3 alone does not restore SA production can be explained by the inhibitory properties of SA on PBS3 activity (*13*). Based on the published crystal structure of PBS3 (*20*), we postulate that SA and isochorismate both occupy the binding pocket, providing the capacity for regulatory feedback inhibition (Fig. S7). In order to overcome competitive inhibition, ectopic overexpression of ICS1 is required to produce sufficient amounts of isochorismate and to allow for quantitative displacement of SA from the active site of PBS3.

By studying SA formation in the autoimmune mutant *snc2-*1D *npr1-*1 *pbs3-*1, we were able to identify isochorismate as the substrate for PBS3 (Fig. 2E). We confirmed that recombinant PBS3 metabolizes isochorismate and glutamate to yield isochorismate-9-glutamate (Fig. 2A). Kinetic analyses (Fig. 2E) as well as *in silico* studies (Fig. S8) corroborated the preference of PBS3 for isochorismate as its native substrate. SA formation via PBS3-derived isochorismate-9-glutamate is about 2 × 10^6^ times faster than its formation via direct chemical decomposition from its precursor isochorismate (*18*). The detection of isochorismate-9-glutamate as well as 2-hydroxy acryloyl-*N*-glutamate *in planta* supported our *in vitro* findings (Fig. 2G and H). Our study suggests that EDS5 exports isochorismate from the plastid into the cytosol, where PBS3 metabolizes it to isochorismate-9-glutamate. PBS3 and its amino acid residues conferring substrate-specific enzymatic activity are highly conserved across the plant kingdom (*26*) (Fig. S8). This substantiates the idea that the PBS3-dependent generation of the unstable intermediate isochorismate-9-glutamate and its spontaneous decomposition represents the canonical mechanism for isochorismate-derived SA biosynthesis in plants.

Shortly after deposition of an earlier version of this manuscript at bioRxiv (*27*), another manuscript was uploaded on the same preprint server (*28*), reporting similar results regarding the role of PBS3 in SA production. However, the authors of this manuscript postulate that completion of SA biosynthesis requires the activity of EPS1. EPS1 encodes a predicted BAHD acyltransferase-family protein exclusively found in the *Brassicaceae* family of higher plants (*28*), and is not transcriptionally co-regulated with *ICS1*, *EDS5*, *PBS3* and many other molecular components that are known to be involved in SA-mediated defense (Fig. S1 and S12), arguing against its role as a major contributor to SA biosynthesis. Irrespective of whether or not EPS1 is rate-limiting for ICS1-dependent SA production in *Arabidopsis* and its relatives, their data strongly support our conclusion that non-enzymatic decomposition of the PBS3-product isochorismate-9-glutamate represents the archetypical mechanism in plants. This irreversible process leads to rapid metabolite channeling and prevents substrate depletion by aromatic amino acid synthesis (*29*). The minimal requirement of only three proteins (ICS1, EDS5 and PBS3), their spatial separation and the unidirectional flux provide an effective strategy to protect the pathway against mutations and pathogenic effectors. In bacteria, SA occurs as an intermediate in siderophore biosynthesis (*30*). These iron chelators are essential for microbial survival and pathogenicity. It is tempting to speculate that the reaction mechanism found here may also occur in human pathogens and thus represents a potential new target for antibiotics (*31*).

## Acknowledgments

We are grateful to Elena Petutschnig (University of Goettingen) for support in confocal microscopy, to Jane Parker (MPIPZ Cologne) for pXCSG-YFP/CFP destination vectors, to Ellen Hornung (University of Goettingen) for pEntry-C-eYFR vector, to Amelie Kelly for critical reading the manuscript and to Egon Fanghänel for advice on chemical decomposition.

## Author contributions

D.R., D.L. Y.D., K.F., V.L., M.W., Y.Z., and I.F. conceived and designed the experiments. D.R., D.L., K.Z. and Y.D. performed the experiments. D.R., D.L., K.F., Y.D., V.L., M.W., Y.Z. and I.F. analyzed and discussed the data, D.R., D.L, K.F., K.Z., Y.Z., M.W., V.L. and I.F. wrote the article.

## Data and materials availability

All data are available in the main text or the supplementary materials. The authors responsible for distribution of materials integral to the findings presented in this article are: Ivo Feussner, Marcel Wiermer and Yuelin Zhang.

## Supplementary Materials

Materials and Methods

Figures S1-S12

Tables S1-S2

References (*1–11*)

