## Supplemental Information for "PBS3 is the missing link in plant-specific isochorismate-derived salicylic acid biosynthesis"

**This PDF file includes:**

Materials and Methods
Figs. S1 to S12
Tables S1 and S2
References (11)

### Materials and Methods

Metabolite fingerprint analysis

The non-targeted metabolite analysis was performed as described in (*1, 2*). In short, metabolites were extracted from ~ 100 mg leaf material or from two-week-old seedlings growing on tissue culture plates by two-phase extraction with methyl-*tert*-butylether (MTBE). Extracts were separated with an Ultra Performance Liquid Chromatography (UPLC, ACQUITY UPLC System; Waters Corporation, USA) equipped with an ACQUITY UPLC HSS T3 column (1.0 x 100 mm, 1.8 µm particle size, Waters Corporation, USA), coupled to a time-of-flight mass spectrometer (TOF-MS, LCT Premier; Waters Corporation, USA).

For peak picking and alignment, MarkerLynx Application Manager 4.1 for MassLynx software was used. For subsequent data processing, ranking, filtering, adduct correction, clustering and database analysis, MarVis software (MarkerVisualization, http://marvis.gobics.de, (*2*)) was utilized. Bars represent the mean ± STD of three biological replicates. Statistical differences among replicates are labeled with different letters ($P < 0.05$, one-way ANOVA and post hoc Tukey's Test; $n = 3$). All experiments were repeated at least twice with similar results.

Accurate mass high resolution MS/MS analysis

The chemical structures of CA, IC, SA, 2HNG, CA-7-Glu, CA-9-Glu, IC-7-Glu and IC-9-Glu were elucidated by MS/MS analyses with an 1290 Infinity UHPLC system coupled to a 6540 UHD Accurate-Mass Q-TOF (Agilent Technologies, USA) as previously described (*2*). Data acquisition was monitored with Mass Hunter Workstation Acquisition software B.05.01 (Agilent Technologies, USA). For data analysis, Mass Hunter Qualitative Analysis software B.05.01 was used (Agilent Technologies, USA). Authentic standards for SA (BioXtra, ≥99.0%, S5922 Sigma, Germany) and CA (≥80%, C1761 Aldrich, Germany) were obtained from Merck, Germany. All fragmentations were carried out in the negative ionization mode. The results are shown in Fig. 3C-E, S3, S5 and are summarized in Table S1.

Quantitative SA and SAG measurements

The absolute amounts of SA and SAG were determined as previously described (*3*).

Protein expression and purification

*AtGH3.12* was amplified by PCR from *A. thaliana* ecotype Col-0 cDNA derived from systemic leaves after infection (*1*). *AtICS1* without the sequence of the transit peptide was amplified from Col-0 cDNA of infected leaves. *AtGH3.12* was subcloned into pET24 (Novagen, Germany), and *AtICS1* into pET28a (Novagen, Germany). Both His-tagged proteins were expressed in *E. coli* Rosetta2 (DE3) and purified with affinity and size-exclusion chromatography. Bacteria were handled as described in (*1*). The cells were disrupted by pulsed ultrasonic with micro-tip (Branson Sonifier Cell Disruptor B15, Branson Ultrasonics Corporation, USA) in a solution containing 100 mM Tris-HCl, pH 7.4, 500 mM NaCl, 1 mM phenylmethylsulfonyl fluoride (PMSF) and 1 mM dithiothreitole (DTT). Cell debris was removed by centrifugation (50000 x g, 4 °C, 30 min). The supernatant was applied onto a HisTrap column (GE Healthcare, GB) pre-equilibrated with 100 mM Tris-HCl, pH 7.4, 500 mM NaCl, 1 mM DTT. For the elution, 30 % elution buffer (100 mM Tris-HCl, pH 7.4, 200 mM NaCl, 500 mM imidazole, 1 mM DTT) was used. Fractions containing the protein of interest were pooled and applied onto a size exclusion chromatography S200 gel filtration column (GE Healthcare, GB), which were pre-equilibrated with 50 mM Tris-HCl, pH 8.0, 10 mM $MgCl_2$, 1 mM DTT. Both chromatographies were operated with an ÄKTAprime plus system (GE Healthcare, GB) at 4°C. The expression levels as well as the purity of the proteins were verified by SDS-PAGE and visualized with Coomassie Brilliant Blue staining. Protein aliquots were stored at -80 °C without noticeable loss of activity.

Activity assays

Isochorismate was synthesized by incubating 0.4 mg/ml His-ICS1 with either 1 or 0.1 mM CA in 100 µL of 50 mM Tris-HCl, pH 7.4 with 5 mM $MgCl_2$, 10 mM NaCl and 1 mM DTT for 10 min at 30 °C and shaking at 150 rpm. The protein was subsequently removed from the solution by ultrafiltration (25°C, 16000 xg) using a SpinX UF 500 concentrator (10,000 MWCO; Corning, USA). The product of the ICS1 reaction was monitored by UHPLC-Q-TOF-MS analyses.

For the qualitative PBS3 activity assay, reactions were performed in 100 µL of 50 mM Tris-HCl, pH 7.8 with 5 mM $MgCl_2$, 4 mM glutamic acid, 5 mM ATP, 0.6 mg/mL His-PBS3 and 6 µM ISC or 50 µM CA. The reaction was performed for 1 h at 30 °C, shaking at 150 rpm and subsequently stopped by addition of 20 µL acetonitrile. The products were monitored by UHPLC-Q-TOF-MS analyses.

Determination of the kinetic constants

To monitor the activity of PBS3 quantitatively, a spectrophotometric assay was employed. The formation of AMP by the PBS3 reaction was coupled to the reactions of pyruvate kinase, myokinase and lactate dehydrogenase (*4*). The assays were performed at 30 °C in 200 µL of 50 mM Tris-HCl, pH 7.8 with 5 mM $MgCl_2$, 50 mM KCl, 0.5 mM phosphoenolpyruvate, 4 mM glutamic acid, 2 mM ATP, 4 units of rabbit muscle pyruvate kinase (P1506, Sigma, Germany), 4 units of rabbit muscle myokinase (M3003, Sigma, Germany) and 4 units of rabbit muscle lactate dehydrogenase (L2500, Sigma, Germany) and 125 µM NADH. The kinetic constants of PBS3 were determined for 4HBA (99%, H20059, Aldrich, Germany) and isochorismate as carboxyl-substrates. Thereby, 0-1000 µM of 4HBA and 50 µg/mL His-PBS3 or 0-35 µM isochorismate and 0.5 µg/mL His-PBS3 were used. The activity of His-PBS3 at lower protein concentration was stabilized with 0.13% (w/v) BSA (8076.2, Roth, Germany). The initial velocity was measured using a V-630 spectrophotometer (Jasco, Germany) by monitoring the absorbance change at 340 nm over 600 s at 30 °C.

Transient expression and subcellular localization

To determine the subcellular localization of PBS3, *PBS3* was sub-cloned from pET-PBS3 into the pENTR vector utilizing the pENTR/D-TOPO Cloning Kit (Invitrogen, USA). The sequence confirmed constructs were used in an LR reaction with the binary destination vector pXCSG-YFP (*5*) to generate *pXCSG 35S::PBS3::YFP*. To determine potential co-localization, full length ICS1 was amplified from cDNA of *Psm.* infected Col-0 leaves (*1*). Full length *ICS1* was ultimately cloned into pXCSG-CFP (4) to generate *pXCSG 35S::ICS1::CFP*. For the construction of a plastidial PBS3, a cDNA fragment encoding the first 47 amino acids of ICS1 were cloned upstream of *PBS3*. The full construct was transformed into the pXCSG-YFP vector to generate *pXCSG 35S::chloroPBS3::YFP*. Additionally, a pUC18-derived pEntry vector containing a C-terminal eYFP coding sequence was utilized to prepare a *pEntry-35S::PBS3::YFP* construct. By an LR reaction, this construct was integrated into the binary pCambia vector to yield *pCambia-35S::PBS3::YFP*. For transient expression in *Arabidopsis* leaves, we followed the protocol from (*6*) but used a higher *Agrobacterium tumefaciens* GV3101 concentration ($OD_{600}$ = 0.4) to enable SA quantification by UHPLC-Q-TOF-MS 24 hours after infiltration.

Confocal laser scanning microscopy was performed on transiently transformed *A. thaliana efr* leaves, 3 days after co-infiltration of *Agrobacteria* carrying the *pXCSG 35S::ICS1::CFP* and *pCambia-35S::PBS3::YFP*, or plastidial *pXCSG 35S::chloroPBS3::YFP* constructs. The *efr* mutant background was used to achieve higher transformation efficiencies to allow fluorescence detection (*7*). Bacterial strains were co-infiltrated with a respective final $OD_{600}$ of 0.3 together with a p19 silencing suppressor. All images for localization were taken with a 20x/0.75 objective (HC PL APO, CS2) of the TSC-SP8 FALCON microscope (Leica, Bensheim, Germany) operated by the LAS X Software (v3.5.1). For fluorescence lifetime imaging (FLIM), images were taken with a 63x/1.20 water immersion objective with correction collar (HC PL APO, CS2). ICS1-CFP was excited using a 440 nm diode laser with a pulse rate of 40 kHz, PBS3-YFP and chloroPBS3-YFP were excited using 514 nm of a white light laser with a pulse rate of 40 kHz. Emission was detected at 454-491 nm for CFP and 525-560 nm for YFP using Leica HyD SDM detectors, while chlorophyll auto-fluorescence was detected at 690-770 nm using a Leica HyD detector. To generate FLIM data, 1000 photons were collected in a 512 x 512 pixel format in the brightest channel for co-infiltrated tissues, while for mock infiltrated tissues the average number of frames obtained for the respective co-infiltration was detected using the same laser and detector settings. The average photon arrival times, as an overlay on intensity for every pixel, were calculated and displayed using the FALCON LASX/FLIM/FCS Software (v3.5.5). Images were sequentially scanned. Merging of images was performed using Fiji software (*8*).

| | LOCUS | NAME | FUNCTION | Supportability | MR |
|---|---|---|---|---|---|
| 0 | At1g74710 | ICS1 | ADC synthase superfamily protein | 3 | 0 |
| 1 | At5g13320 | PBS3 | Auxin-responsive GH3 family protein | 3 | 2.8 |
| 2 | At4g39030 | EDS5 | MATE efflux family protein | 3 | 10 |
| 3 | At3g47480 | Calcium-binding EF-hand family protein | Calcium-binding EF-hand family protein | 3 | 16.3 |
| 4 | At1g33960 | IAN8 | P-loop containing nucleoside triphosphate hydrolases superfamily protein | 3 | 19 |
| 5 | At2g04450 | NUDX6 | nudix hydrolase homolog 6 | 3 | 19 |
| 6 | At4g23150 | CRK7 | cysteine-rich RLK (RECEPTOR-like protein kinase) 7 | 3 | 27 |
| 7 | At2g18660 | PNP-A | plant natriuretic peptide A | 3 | 27.1 |
| 8 | At3g48090 | EDS1 | alpha/beta-Hydrolases superfamily protein | 3 | 28 |
| 9 | At2g46400 | WRKY46 | WRKY DNA-binding protein 46 | 3 | 28.1 |
| 10 | At3g52430 | PAD4 | alpha/beta-Hydrolases superfamily protein | 3 | 30.4 |
| 11 | At5g52760 | Copper transport | Copper transport protein family | 3 | 30.8 |
| 12 | At1g21240 | WAK3 | wall associated kinase 3 | 3 | 31.2 |
| 13 | At4g11890 | ARCK1 | Protein kinase superfamily protein | 3 | 33.6 |
| 14 | At1g19250 | FMO1 | flavin-dependent monooxygenase 1 | 3 | 36.9 |
| 15 | At3g48640 | Transmembrane protein | Transmembrane protein | 3 | 42.4 |
| 16 | At5g39670 | Calcium-binding EF-hand family protein | Calcium-binding EF-hand family protein | 3 | 42.5 |
| 17 | At4g38560 | Arabidopsis phospholipase-like protein (PEARLI 4) family | Arabidopsis phospholipase-like protein (PEARLI 4) family | 3 | 44.5 |
| 18 | At4g14365 | XBAT34 | XB3 ortholog 4 in Arabidopsis thaliana | 3 | 46.3 |
| 19 | At4g04490 | CRK36 | cysteine-rich RLK (RECEPTOR-like protein kinase) 36 | 3 | 46.3 |
| 20 | At3g57260 | PR2 | beta-1,3-glucanase 2 | 3 | 49 |
| 21 | At4g04500 | CRK37 | cysteine-rich RLK (RECEPTOR-like protein kinase) 37 | 3 | 49.1 |
| 22 | At1g13470 | Protein of unknown function (DUF1262) | Protein of unknown function (DUF1262) | 3 | 51.8 |
| 23 | At3g13100 | MRP7 | multidrug resistance-associated protein 7 | 3 | 52.6 |
| 24 | At5g64000 | SAL2 | Inositol monophosphatase family protein | 3 | 54.7 |
| 25 | At4g23610 | LEA | Late embryogenesis abundant (LEA) hydroxyproline-rich glycoprotein family | 3 | 54.8 |
| 26 | At2g37710 | RLK | receptor lectin kinase | 3 | 59.5 |
| 27 | At1g66880 | Protein kinase superfamily protein | Protein kinase superfamily protein | 3 | 61.6 |
| 28 | At4g03450 | Ankyrin repeat family protein | Ankyrin repeat family protein | 3 | 63.5 |
| 29 | At1g73805 | SARD1 | Calmodulin binding protein-like | 3 | 63.6 |
| 30 | At1g08450 | CRT3 | calreticulin 3 | 3 | 65.6 |
| 31 | At1g09080 | BIP3 | Heat shock protein 70 (Hsp 70) family protein | 3 | 66.4 |
| 32 | At1g57630 | Toll-Interleukin-Resistance (TIR) domain family protein | Toll-Interleukin-Resistance (TIR) domain family protein | 3 | 66.4 |
| 33 | At1g35230 | AGP5 | arabinogalactan protein 5 | 3 | 66.7 |
| 34 | At2g32680 | RLP23 | receptor like protein 23 | 3 | 67.3 |
| 35 | At3g47540 | Chitinase family protein | Chitinase family protein | 3 | 70.5 |
| 36 | At5g10760 | AED1 | Eukaryotic aspartyl protease family protein | 3 | 72.5 |
| 37 | At5g26920 | CBP60G | Cam-binding protein 60-like G | 3 | 72.9 |
| 38 | At3g60420 | Phosphoglycerate mutase family protein | Phosphoglycerate mutase family protein | 2 | 76.8 |
| 39 | At4g23210 | CRK13 | cysteine-rich RLK (RECEPTOR-like protein kinase) 13 | 3 | 78.4 |
| 40 | At2g14610 | PR1 | pathogenesis-related gene 1 | 3 | 78.4 |
| 41 | At2g24850 | TAT3 | tyrosine aminotransferase 3 | 3 | 87.7 |
| 42 | At5g17990 | TRP1 | tryptophan biosynthesis 1 | 3 | 95.7 |
| 43 | At2g19190 | FRK1 | FLG22-induced receptor-like kinase 1 | 3 | 96.6 |
| 44 | At3g25010 | RLP41 | receptor like protein 41 | 3 | 98.8 |
| 45 | At1g75040 | PR5 | pathogenesis-related gene 5 | 3 | 100.2 |
| 46 | At1g35710 | Protein kinase with LRR domain | Protein kinase family protein with leucine-rich repeat domain | 3 | 100.8 |
| 47 | At3g28540 | P-loop containing nucleoside triphosphate hydrolases superfamily protein | P-loop containing nucleoside triphosphate hydrolases superfamily protein | 3 | 101 |
| 48 | At2g17040 | NAC036 | NAC domain containing protein 36 | 3 | 101.4 |
| 49 | At5g24210 | alpha/beta-Hydrolases | alpha/beta-Hydrolases superfamily protein | 3 | 101.4 |
| 50 | At1g21250 | WAK1 | cell wall-associated kinase | 3 | 104.3 |
| 51 | At1g10340 | Ankyrin repeat family protein | Ankyrin repeat family protein | 3 | 106.7 |
| 52 | At1g44130 | Eukaryotic aspartyl protease family protein | Eukaryotic aspartyl protease family protein | 3 | 107.6 |
| 53 | At5g52810 | SARD4 | NAD(P)-binding Rossmann-fold superfamily protein | 3 | 110 |
| 54 | At3g01830 | Calcium-binding EF-hand family protein | Calcium-binding EF-hand family protein | 3 | 111.4 |
| 55 | At5g03350 | LLP1 | Legume lectin family protein | 3 | 112.4 |
| 56 | At2g20142 | Toll-Interleukin-Resistance (TIR) domain family protein | Toll-Interleukin-Resistance (TIR) domain family protein | 3 | 112.7 |
| 57 | At3g13950 | Ankyrin repeat family protein | | 3 | 113.6 |
| 58 | At2g31880 | SOBIR1 | Leucine-rich repeat protein kinase family protein | 3 | 115.6 |
| 59 | At4g33050 | IQM1 | calmodulin-binding family protein | 3 | 117.8 |
| 60 | At5g01540 | LECRK-VI.2 | lectin receptor kinase a4.1 | 3 | 119 |
| 61 | At5g61010 | EXO70E2 | exocyst subunit exo70 family protein E2 | 3 | 122.5 |
| 62 | At1g01560 | MPK11 | MAP kinase 11 | 3 | 124.3 |
| 63 | At1g76040 | CPK29 | calcium-dependent protein kinase 29 | 3 | 126.5 |
| 64 | At5g19240 | Glycoprotein membrane precursor GPI-anchored | Glycoprotein membrane precursor GPI-anchored | 3 | 127.3 |
| 65 | At5g10380 | RING1 | RING/U-box superfamily protein | 3 | 130.8 |
| 66 | At3g28510 | P-loop containing nucleoside triphosphate hydrolases superfamily protein | P-loop containing nucleoside triphosphate hydrolases superfamily protein | 3 | 131.6 |
| 67 | At2g04430 | NUDT5 | nudix hydrolase homolog 5 | 3 | 132.4 |
| 68 | At2g29120 | GLR2.7 | glutamate receptor 2.7 | 3 | 133.9 |
| 69 | At1g78410 | VQ motif | VQ motif-containing protein | 3 | 135.1 |
| 70 | At4g10500 | DLO1 | 2-oxoglutarate (2OG) and Fe(II)-dependent oxygenase superfamily protein | 3 | 137.9 |

**Fig. S1.** Expression of *EDS5* (At4g39030) and *PBS3* (At5g13320) is co-regulated with *ICS1* (At1g74710). Co-expression analysis was performed using *ICS1* as bait (marked red) with

ATTED-II (http://atted.jp/; Version: 9.2). The top 70 ranked co-expressed genes from microarray-based datasets are displayed. *EDS5* and *PBS3* are marked in green, immunity related genes are marked in yellow. Supportability is displayed by numbers (0-3) according to the p-value threshold (0=>1E-01, 1 =<1E-01, 2=<1E-01, 3=<1E-03) as a measure of reproducibility. The mutual rank (MR) index is indicated as a measure of co-expression (*9*).

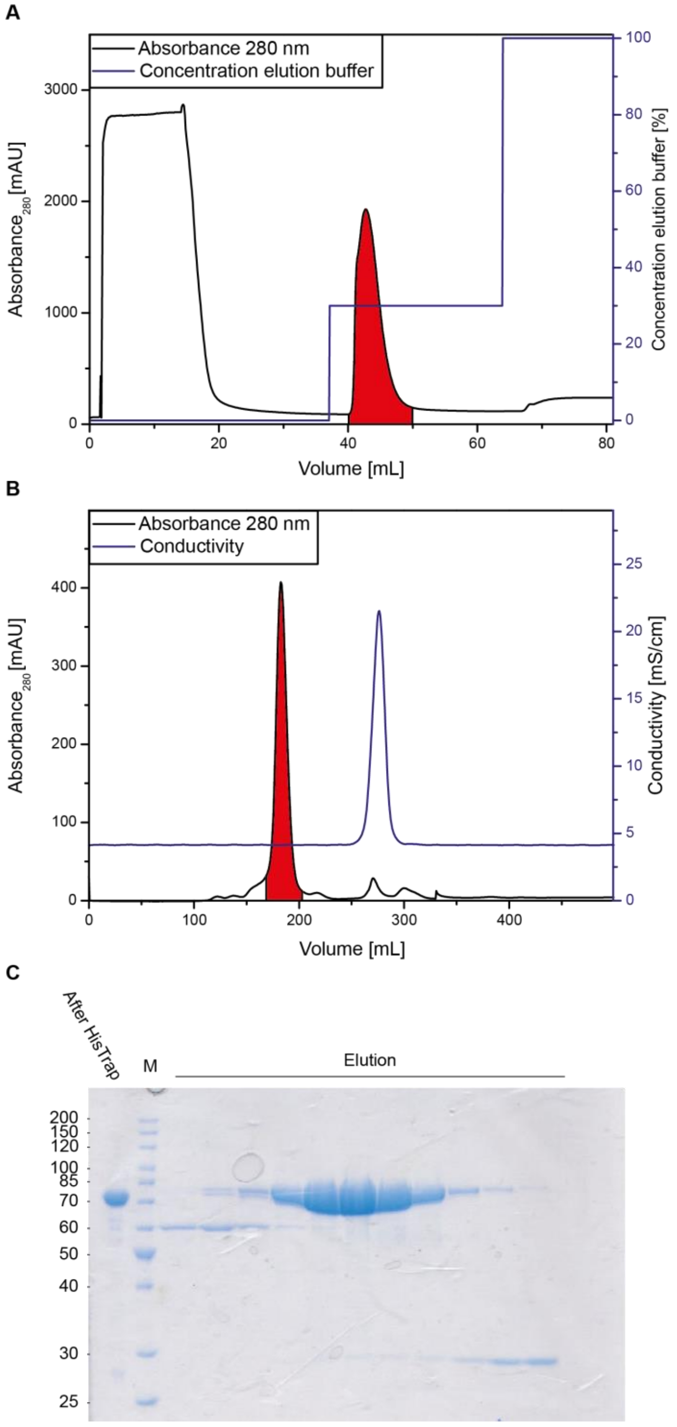


**Fig. S2.** Protein purification of His6-tagged AtPBS3 via affinity - and size exclusion chromatography. PBS3 was heterologously expressed in *E. coli* and subsequently purified via affinity chromatography (A). Fractions corresponding to the area marked in red were collected and applied to size exclusion chromatography (B). Fractions containing proteins were collected and examined by SDS-PAGE (C) to verify the purity. Pure protein containing fractions with the predicted molecular mass were pooled and concentrated by filter centrifugation.

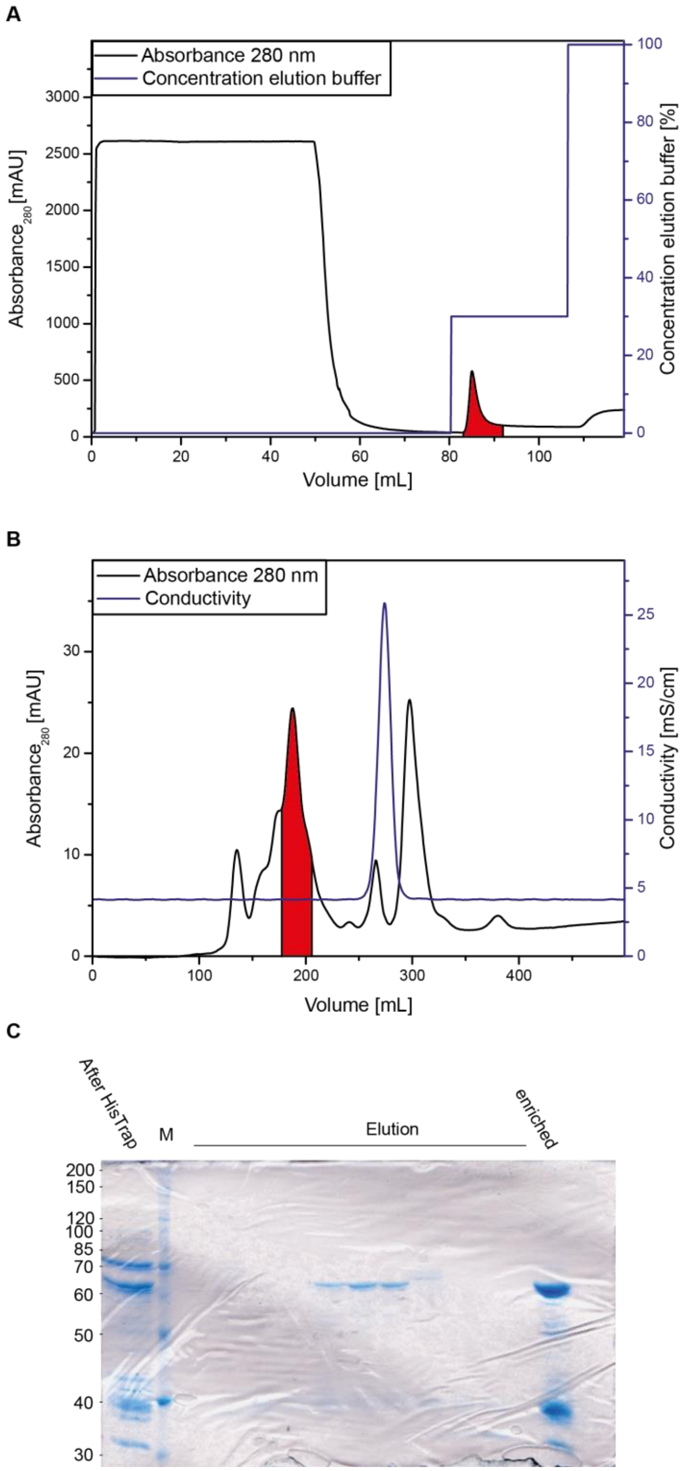


**Fig. S3.** Protein purification of His6-tagged AtICS1 via affinity - and size exclusion chromatography. ICS1 was heterologously expressed in *E. coli* and subsequently purified via affinity chromatography (A). Fractions corresponding to the area marked in red were collected and applied to size exclusion chromatography (B). Fractions containing proteins were collected and examined by SDS-PAGE (C) to verify the purity. Pure protein containing fractions with the predicted molecular mass were pooled and concentrated by filter centrifugation.

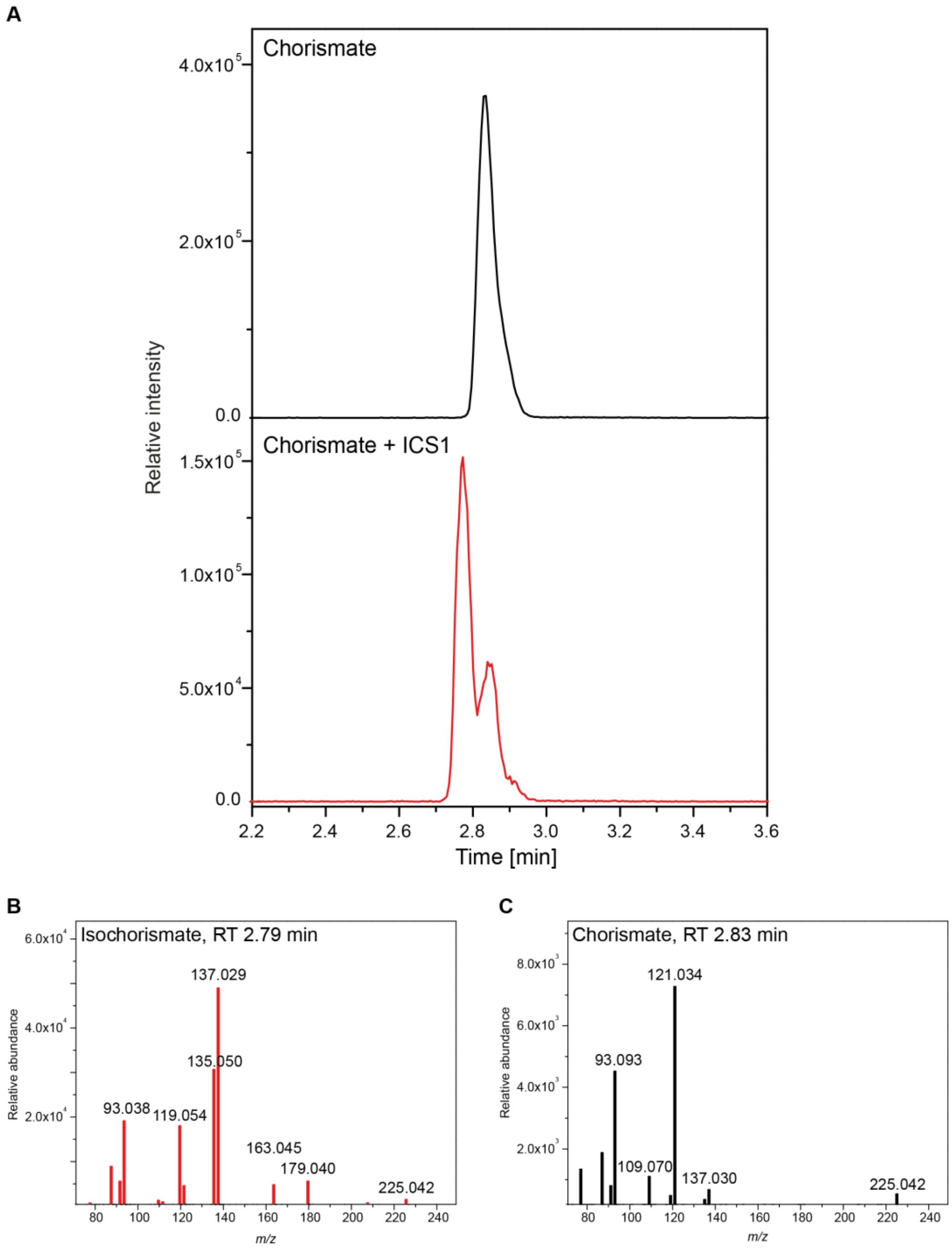


**Fig. S4.** Incubation of ICS1 with chorismate yields isochorismate. LC-MS analysis was performed on activity assays of purified ICS1 with chorismate. (A) The extracted ion chromatogram for chorismate/isochorismate (*m/z* 225.042) is shown. Upper panel without -, lower panel with enzyme. (B) MS/MS fragmentation for isochorismate from the incubation mixture at 2.79 min. (C) MS/MS fragmentation for chorismate from the activity assay without enzyme at 2.83 min. Isochorismate and chorismate can be distinguished based on retention time shift and differences in their fragmentation pattern. RT: retention time.

A

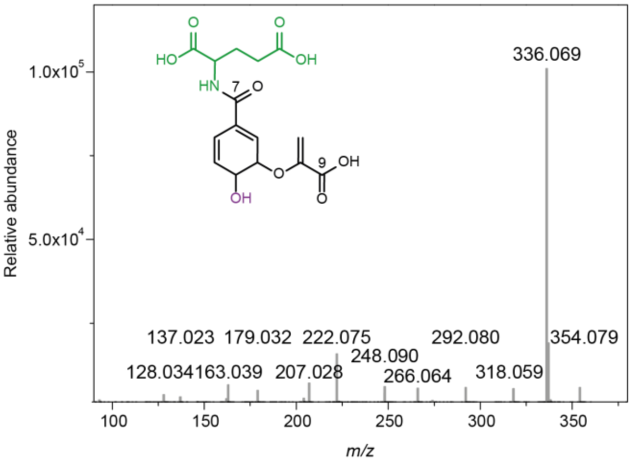


B

| MS² fragment of CA-7-Glu [*m/z*] | Elemental composition | Identity |
|---|---|---|
| 354.079 | $C_{15}H_{16}NO_9$ | $[M\text{-}H]^-$ |
| 336.069 | $C_{15}H_{14}NO_8$ | $[M\text{-}H_2O\text{-}H]^-$ |
| 318.059 | $C_{15}H_{13}NO_7$ | $[M\text{-}2H_2O]^-$ |
| 292.080 | $C_{14}H_{14}NO_6$ | $[M\text{-}H_2O\text{-}CO_2\text{-}H]^-$ |
| 266.064 | $C_{12}H_{12}NO_6$ | $[M\text{-}C_3H_4O_3\text{-}H]^-$ |
| 248.090 | $C_{13}H_{14}NO_4$ | $[M\text{-}H_2O\text{-}2CO_2\text{-}H]^-$ |
| 222.075 | $C_{11}H_{12}NO_4$ | $[M\text{-}C_3H_4O_3\text{-}CO_2\text{-}H]^-$ |
| 207.028 | $C_{10}H_7O_5$ | $[M\text{-}Glu\text{-}H]^-$ |
| 179.032 | $C_9H_7O_4$ | $[M\text{-}Glu\text{-}CO\text{-}H]^-$ |
| 163.039 | $C_9H_7O_3$ | $[M\text{-}Glu\text{-}CO_2\text{-}H]^-$ |
| 137.023 | $C_7H_5O_3$ | $[M\text{-}Glu\text{-}C_3H_3O_2]^-$ |
| 128.034 | $C_5H_6NO_3$ | $[Glu\text{-}H_2O\text{-}H]^-$ |

C

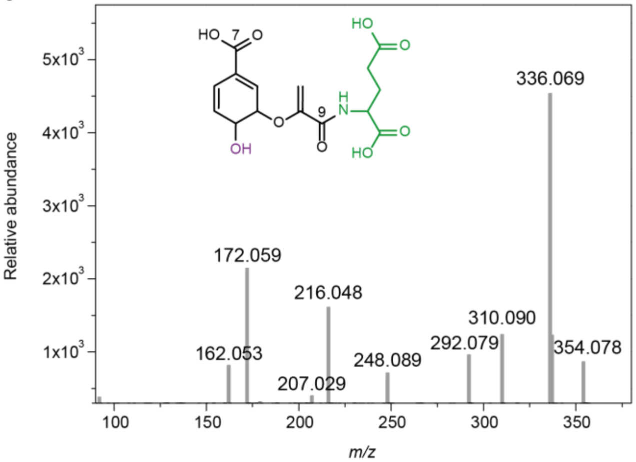


D

| MS² fragment of CA-9-Glu [*m/z*] | Elemental composition | Identity |
|---|---|---|
| 354.078 | $C_{15}H_{16}NO_9$ | $[M\text{-}H]^-$ |
| 336.069 | $C_{15}H_{14}NO_8$ | $[M\text{-}H_2O\text{-}H]^-$ |
| 310.090 | $C_{14}H_{16}NO_7$ | $[M\text{-}CO_2\text{-}H]^-$ |
| 292.079 | $C_{14}H_{14}NO_6$ | $[M\text{-}H_2O\text{-}CO_2\text{-}H]^-$ |
| 248.089 | $C_{13}H_{14}NO_4$ | $[M\text{-}H_2O\text{-}2CO_2\text{-}H]^-$ |
| 207.029 | $C_{10}H_7O_5$ | $[M\text{-}Glu\text{-}H]^-$ |
| 216.048 | $C_8H_{10}NO_6$ | $[M\text{-}SA\text{-}H]^-$ |
| 172.059 | $C_7H_{10}NO_4$ | $[M\text{-}SA\text{-}CO_2\text{-}H]^-$ |
| 162.053 | $C_9H_8NO_2$ | $[M\text{-}CO_2\text{-}H_2O\text{-}C_5H_6O_4\text{-}H]^-$ |

E

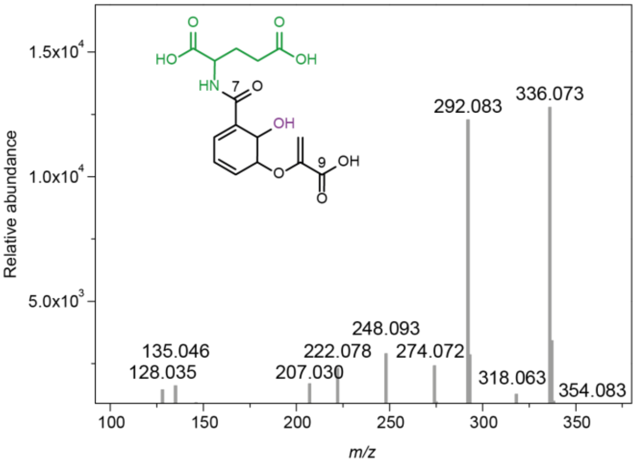


F

| MS² fragment of IC-7-Glu [*m/z*] | Elemental composition | Identity |
|---|---|---|
| 354.083 | $C_{15}H_{16}NO_9$ | $[M\text{-}H]^-$ |
| 336.073 | $C_{15}H_{14}NO_8$ | $[M\text{-}H_2O\text{-}H]^-$ |
| 318.063 | $C_{15}H_{13}NO_7$ | $[M\text{-}2H_2O]^-$ |
| 292.083 | $C_{14}H_{14}NO_6$ | $[M\text{-}H_2O\text{-}CO_2\text{-}H]^-$ |
| 274.072 | $C_{14}H_{12}NO_5$ | $[M\text{-}2H_2O\text{-}CO_2\text{-}H]^-$ |
| 248.093 | $C_{13}H_{14}NO_4$ | $[M\text{-}H_2O\text{-}2CO_2\text{-}H]^-$ |
| 222.078 | $C_{11}H_{12}NO_4$ | $[M\text{-}C_3H_4O_3\text{-}CO_2\text{-}H]^-$ |
| 207.030 | $C_{10}H_7O_5$ | $[M\text{-}Glu\text{-}H]^-$ |
| 135.046 | $C_8H_7O_2$ | $[M\text{-}Glu\text{-}CO\text{-}CO_2\text{-}H]^-$ |
| 128.035 | $C_5H_6NO_3$ | $[Glu\text{-}H_2O\text{-}H]^-$ |

**Fig. S5.** MS/MS fragmentations of the PBS3 *in vitro* products chorismate-7-glutamate (CA-7-Glu), chorismate-9-glutamate (CA-9-Glu) and isochorismate-7-glutamate (IC-7-Glu). PBS3 activity assay was performed with glutamate, ATP and chorismate (A, C) or isochorismate (E) as substrate. MS/MS fragmentation patterns of CA-7-Glu (A), CA-9-Glu (C) and IC-7-Glu (E) with the corresponding annotation lists for the depicted fragments (B, D, F). The identification is based on an accurate mass analysis.

A

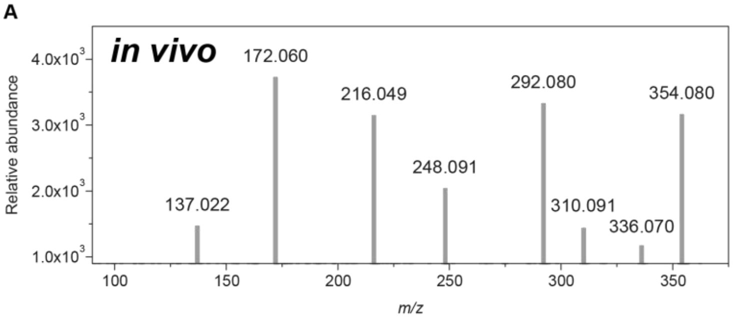


C

| MS² fragment of IC-9-Glu [*m/z*] | Elemental composition | Identity |
|---|---|---|
| 354.080 | $C_{15}H_{16}NO_9$ | $[M-H]^-$ |
| 336.070 | $C_{15}H_{14}NO_8$ | $[M-H_2O-H]^-$ |
| 310.091 | $C_{14}H_{16}NO_7$ | $[M-CO_2-H]^-$ |
| 292.080 | $C_{14}H_{14}NO_6$ | $[M-H_2O-CO_2-H]^-$ |
| 248.091 | $C_{13}H_{14}NO_4$ | $[M-H_2O-2CO_2-H]^-$ |
| 216.049 | $C_8H_{10}NO_6$ | $[M-SA-H]^-$ |
| 172.060 | $C_7H_{10}NO_4$ | $[M-SA-CO_2-H]^-$ |
| 137.022 | $C_7H_5O_3$ | $[SA-H]^-$ |

B

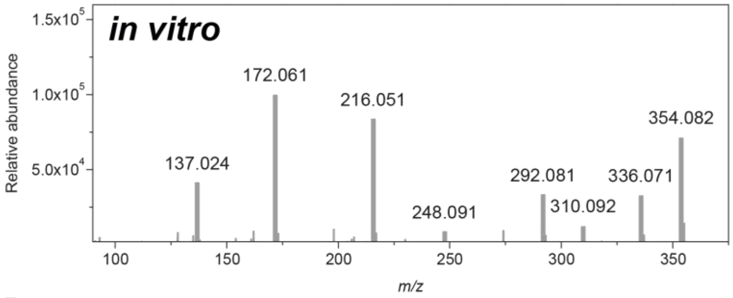


D

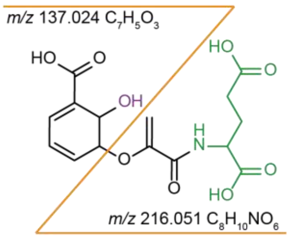


E

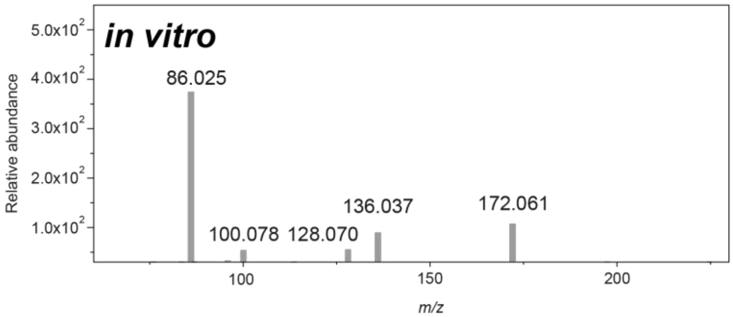


F

| MS² fragment of 2HNG[*m/z*] | Elemental composition | Identity |
|---|---|---|
| 216.051 | $C_8H_{10}NO_6$ | $[M-H]^-$ |
| 172.061 | $C_7H_{10}NO_4$ | $[M-CO_2-H]^-$ |
| 136.037 | $C_7H_6NO_2$ | $[M-CO_2-2H_2O]^-$ |
| 128.070 | $C_6H_{10}NO_2$ | $[M-2CO_2-H]^-$ |
| 100.078 | $C_5H_{10}NO$ | $[M-COH-2CO_2]^-$ |
| 86.025 | $C_3H_4NO_2$ | $[M-C_5H_5O_4-H]^-$ |

**Fig. S6.** MS/MS fragmentations of PBS3 *in vitro* and *in vivo* products. MS/MS fragmentation of isochorismate-9-glutamate (IC-9-Glu) from *snc2*-1D *npr1*-1 plant material (A) and PBS3 activity assay (B). (C) Annotation list for MS/MS fragments of IC-9-Glu from *snc2*-1D *npr1*-1 plant material. (D) Fragment scheme of IC-9-Glu. (E) MS/MS fragmentation of 2-hydroxy acryloyl-N-glutamic acid (2HNG) from the PBS3 activity assay. (F) Annotation list for MS/MS fragments of 2HNG.

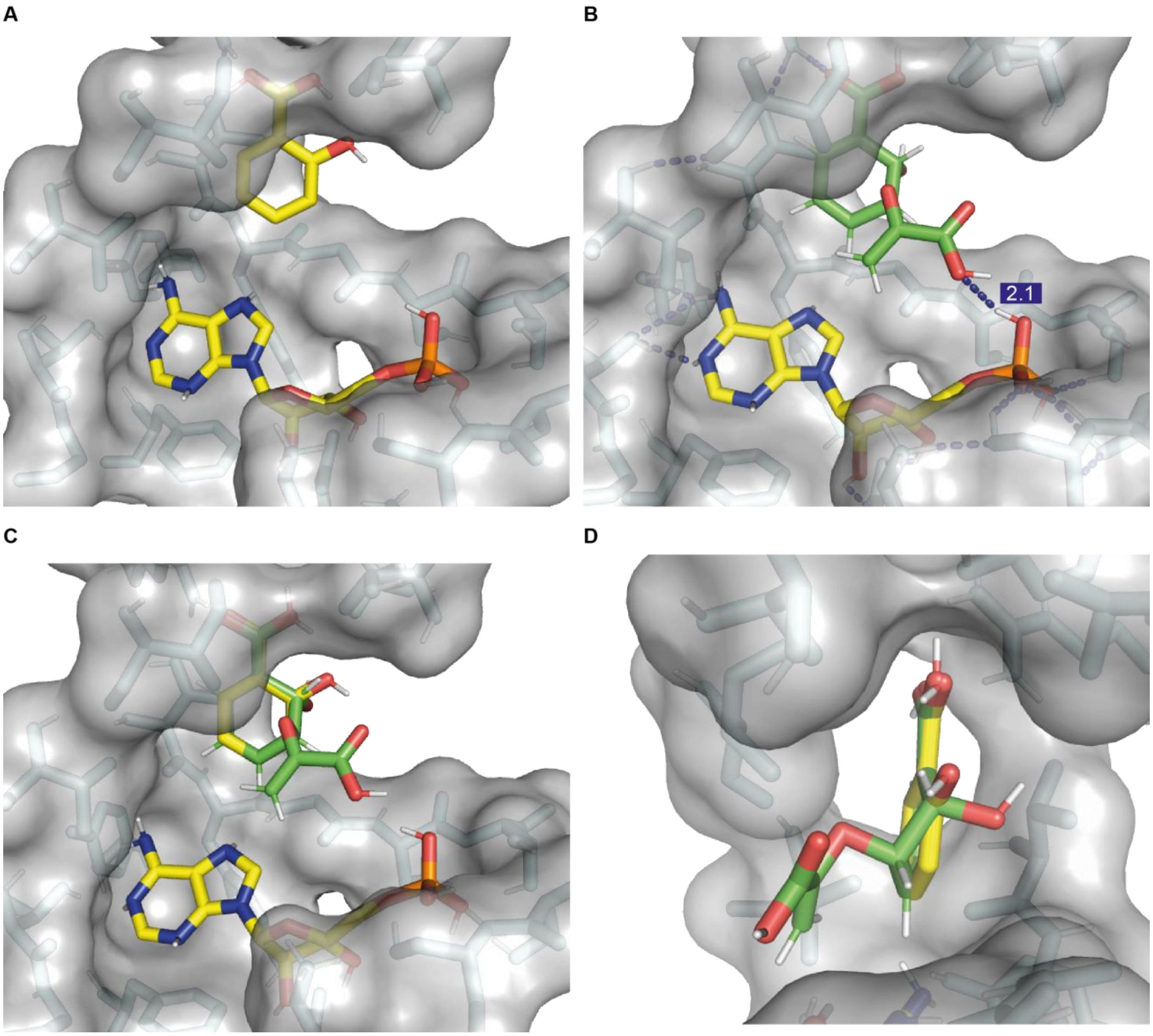


**Fig. S7.** Isochorismate fits into the active site of PBS3. For substrate modeling, the crystal structure of PBS3-AMP-SA (PDB ID 4eql) was used (*10*). Residues in a radius of 4 Å are shown as gray surface to visualize the binding pocket of AMP and the carboxyl substrate. C-atoms of AMP and SA are shown in yellow and for isochorismate in green. **(A)** The large distance between the carboxyl group of SA and the phosphate group of AMP was previously proposed to be the reason why SA is a poor substrate of PBS3 (*10*). **(B)** The hydroxy acrylic group of isochorismate points towards the phosphate group of AMP. The presented model is based on the SA position in the crystal structure and shows the proximity between the carboxyl- and the phosphate group (~2.1 Å). **(C, D)** Overlay of SA (yellow) and isochorismate (green) in the binding pocket of PBS3. Although isochorismate is slightly larger than SA, there appears to be no negative interference with any surrounding residues. This figure was generated with PyMOL (version 2.2.0, Schrödinger, USA).

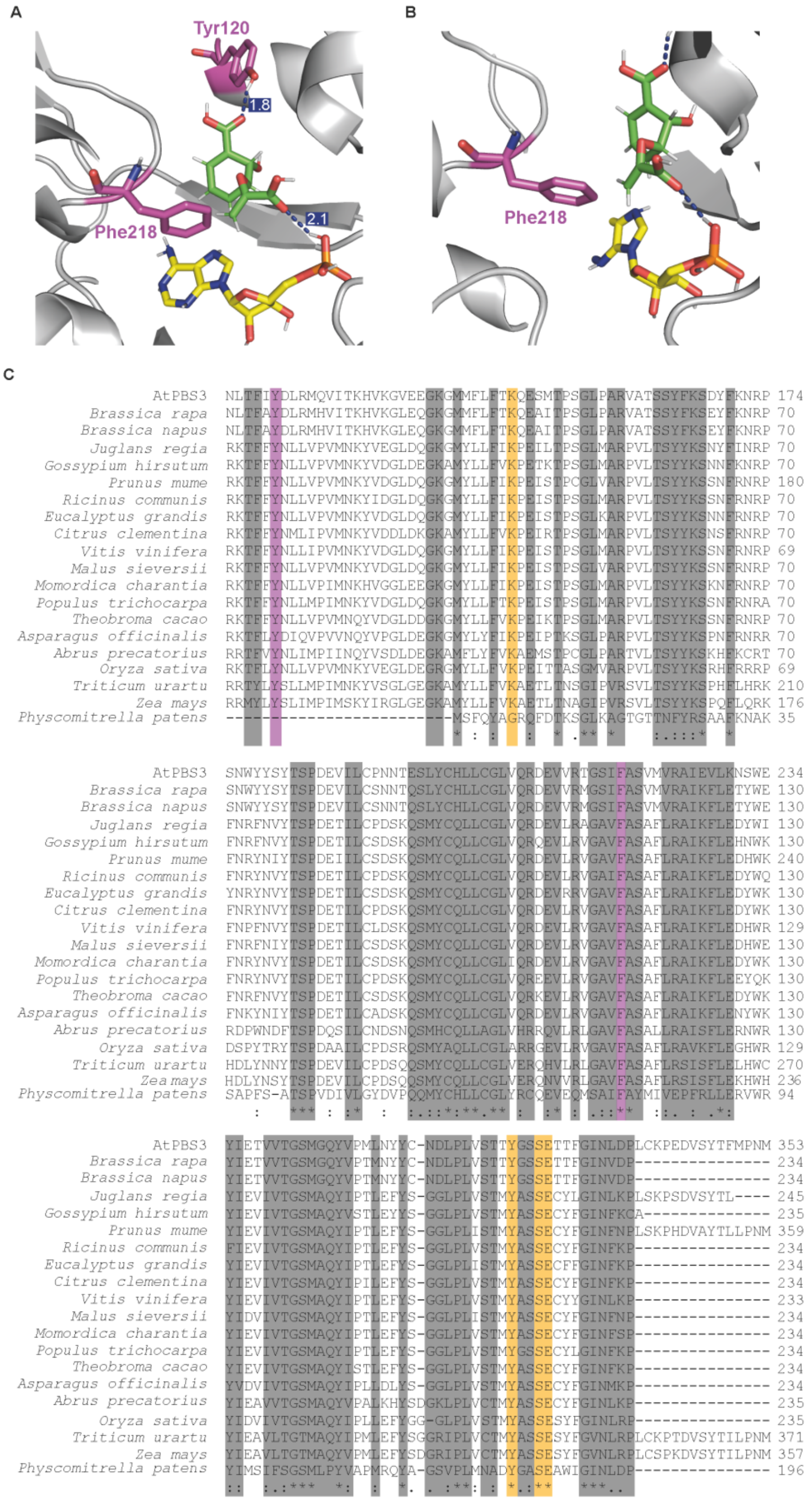


**Fig. S8.** Sequence-based alignment of PBS3 homologs among the plant kingdom. (A, B) Utilizing the crystal structure of PBS3-AMP-SA (PDB ID 4eql) (*10*) with docked isochorismate instead of SA with the binding pocket. Two highly conserved residues of the acyl acid binding site tyrosine120 (Tyr120) and phenylalanine218 (Phe218) are depicted in magenta. Tyr120 interacts with the carboxy group of isochorismate, whereas Phe218 positions the pyrovyl group towards AMP. In this configuration, the distance between the carboxy group of the ring and Tyr120 is ~1.8 Å and between the carboxy group of the pyrovyl group and AMP ~2.1 Å. Both residues are highly conserved within the plant kingdom, as shown in the sequence alignment (C). While Phe218 is present among all studied species, Tyr120 is absent in *Physcomitrella patens* (both residues are highlighted in magenta). This may be due to incomplete sequence annotation of *P. patens*. Highly conserved residues are marked in grey. Residues previously shown to be important for substrate binding (*10*) are highlighted in orange. The comparison aims to compare a wide range of species. Shown here are the following sequences: *Brassica rapa* XP_009125902.1, *Brassica napus* XP_013679572.2, *Juglans regia* XP_018829092.1, *Gossypium hirsutum* XP_016692536.1, *Prunus mume* XP_008222270.1, *Ricinus communis* XP_015570432.1, *Eucalyptus grandis* XP_010066475.1, *Citrus clementina* XP_024040137.1, *Vitis vinifera* XM_002268242.3, *Malus sieversii* AFG33002.1, *Momordica charantia* XP_022153513.1, *Populus trichocarpa* XP_024452922.1, *Theobroma cacao* XP_017972080.1, *Asparagus officinalis* XP_020247065.1, *Abrus precatorius* XP_027367957.1, *Oryza sativa* XM_015760301.2, *Triticum urartu* EMS61902.1, *Zea mays* XP_008653453, *Physcomitrella patens* XP_024386895.1. Only sequence sections containing known substrate binding residues are depicted. Sequence aligment was generated using Clustal Omega (*11*).

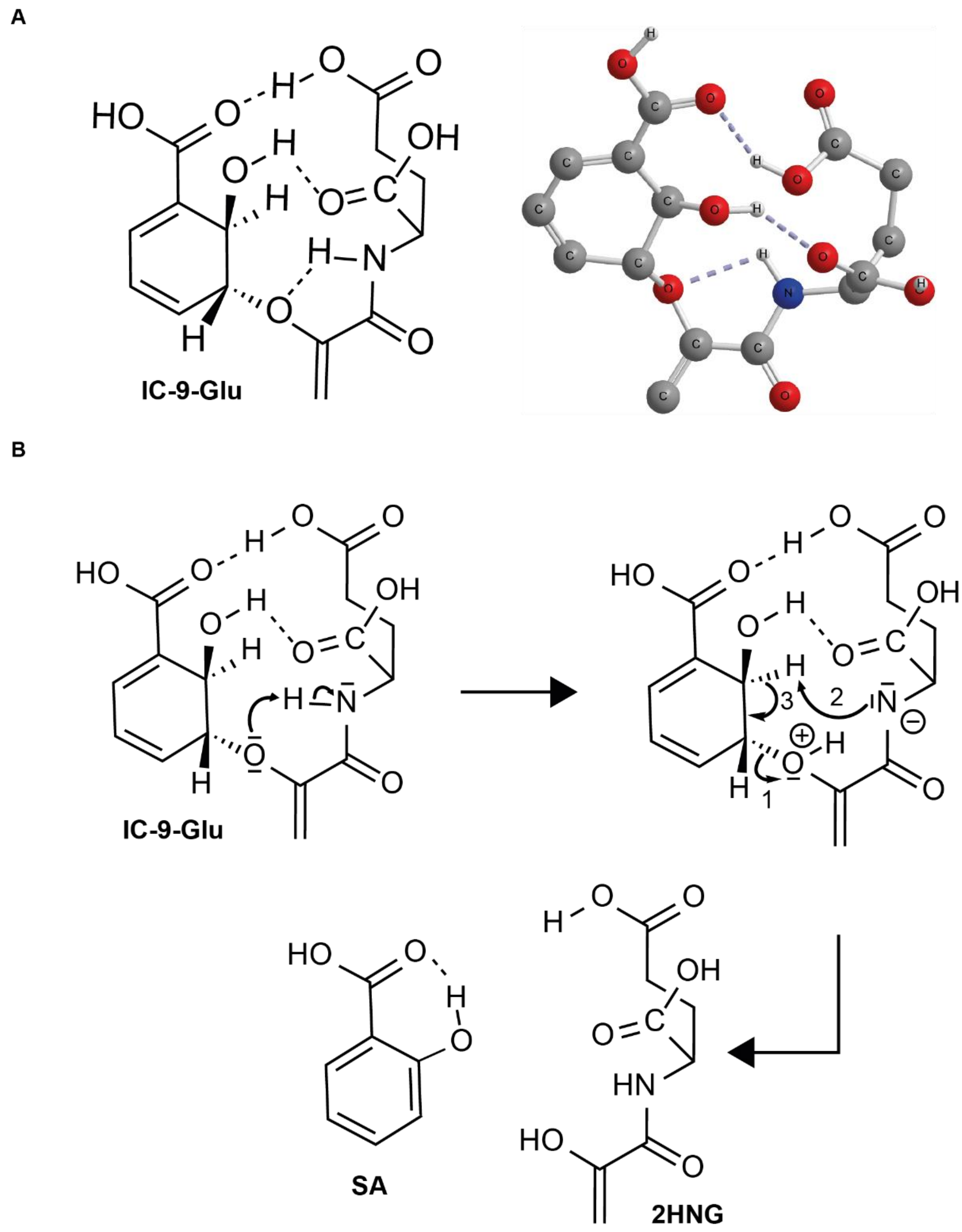


**Fig. S9.** Molecular model of isochorismate-9-glutamate (IC-9-Glu) and its decomposition to SA and 2-hydroxy acryloyl-*N*-glutamate (2HNG). (A) Molecular modeling was used to generate a 3D model of IC-9-Glu**.** This figure was generated with Chem3D (version 17.1 PerkinElmer, USA). (B) The formation of the intramolecular hydrogen bonds facilitates the hydrogen transfer from the amide hydrogen onto the ether oxygen. Subsequently, the non-enzymatic decomposition of IC-9-Glu probably follows an E1 elimination mechanism. In the first part of the reaction, a heterolytic C-O cleavage occurs, which is enhanced by the previous protonation of the linker-oxygen. In the second part, we propose a base initiated aromatization of the ring structure. The final products of this elimination are SA and 2HNG.

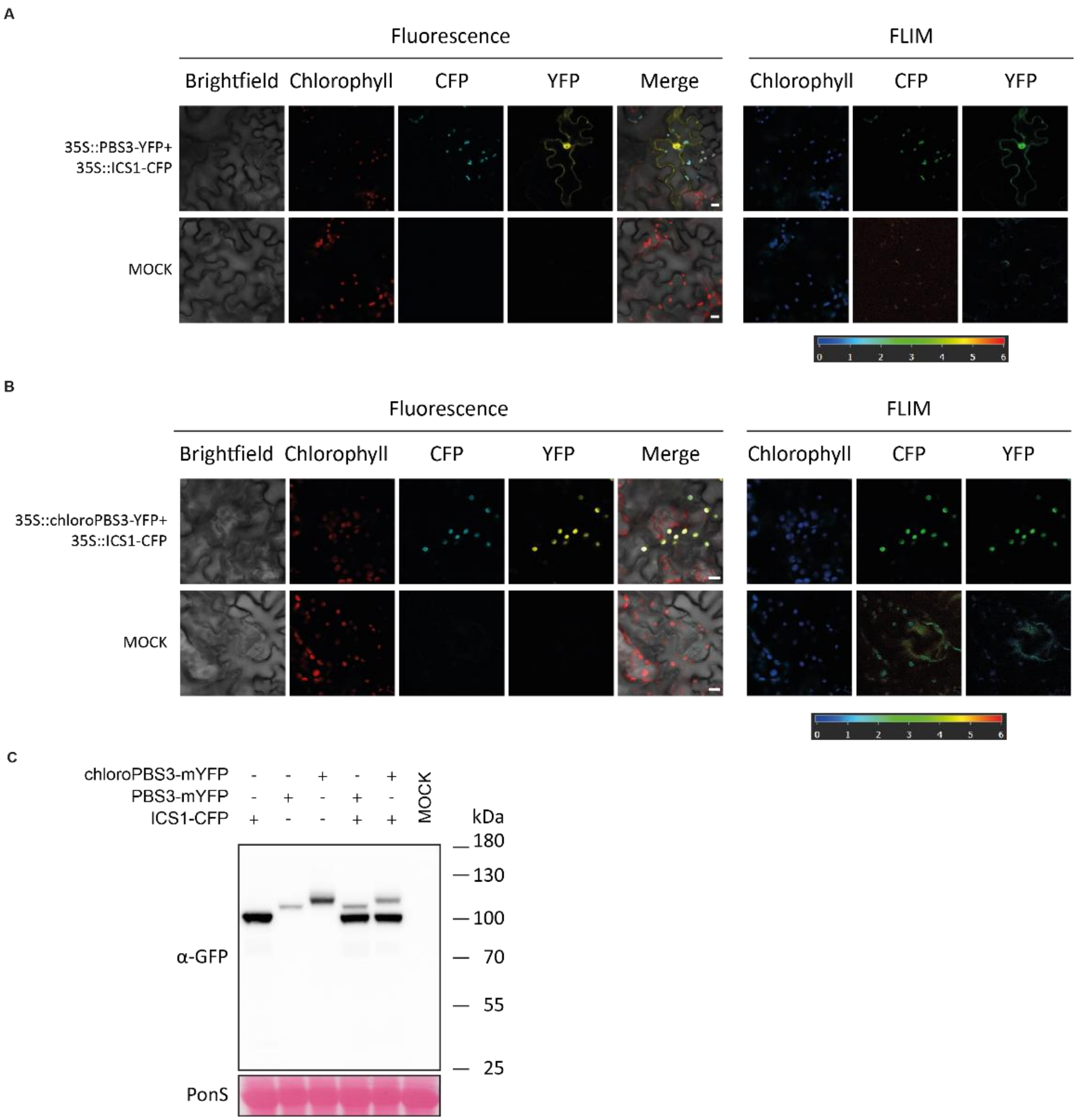


**Fig. S10.** Fluorescence intensity and lifetime images of transiently transformed *Arabidopsis efr* leaf tissue. Fluorescence intensity and average fluorescence lifetime imaging (FLIM) was performed three days after *Agrobacterium*-infiltration for transient co-expression of ICS1-CFP with PBS3-YFP (A) or ICS1-CFP with chloroPBS3-YFP (B) under control of the *35S* promoter in *Arabidopsis efr*. The average lifetime of the detected fluorescence signal is displayed by the hue corresponding to the depicted scale (0 to 6 ns). All constructs were co-expressed together with a p19 silencing suppressor. The same laser and detector settings were applied to analyze mock (p19 only) infiltrated tissues. The scale bar in the merged image of the fluorescence intensity represents 10 µm. (C) Immunoblot analyses of 60 µg total protein extracts of *Arabidopsis efr* infiltrated with *Agrobacteria* for transient co-expression of the indicated fusion proteins. Total extracts were taken three days after *Agrobacterium*-infiltration and were separated on 7.5 % SDS polyacrylamide gels, blotted onto nitrocellulose membranes and probed with α-GFP primary antibody and (α-mouse) secondary antibody coupled to horseradish peroxidase. Loading was monitored by PonceauS (PonS) staining of the membrane.

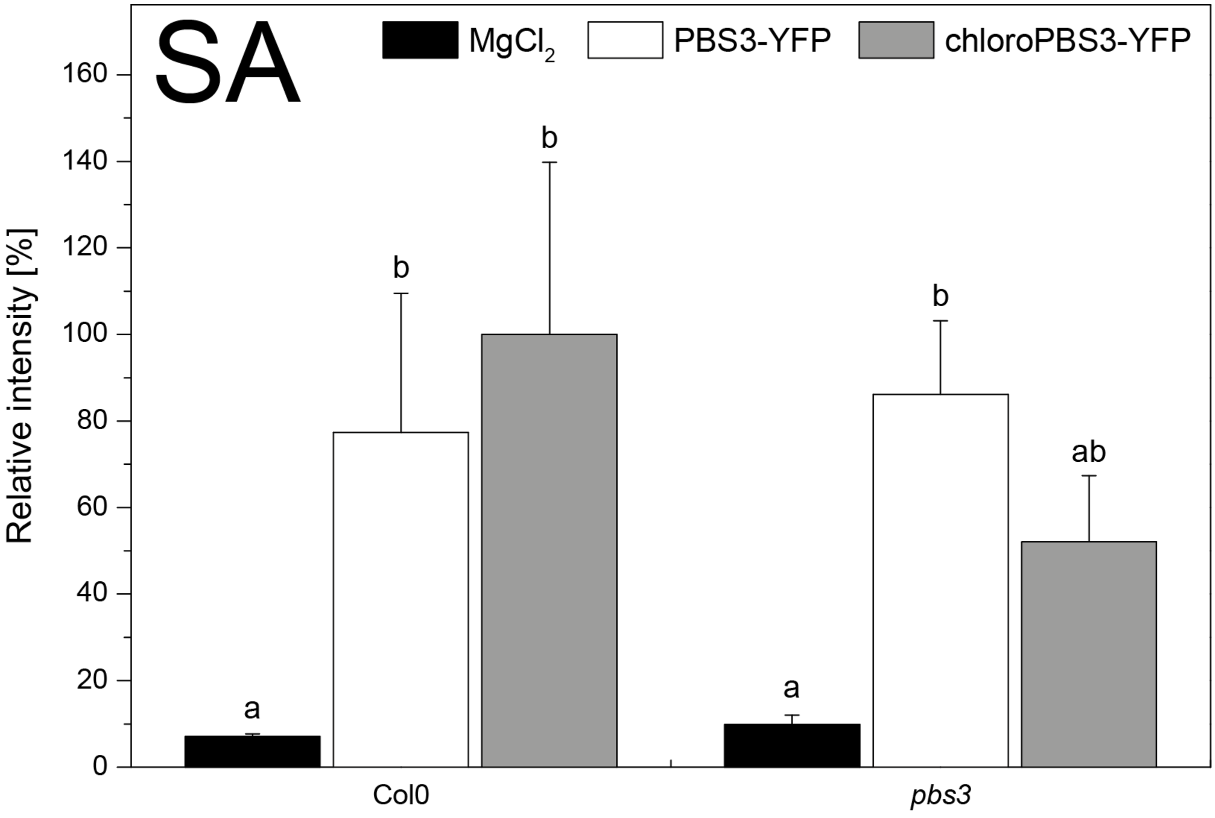


**Fig. S11.** Amounts of *Agrobacterium*-induced salicylic acid (SA) in Col0 and *pbs3-1*. PBS3-YFP or chloroPBS3-YFP was transiently expressed in *Arabidopsis* Col-0 or *pbs3-1* leaves to determine whether they can restore the *Agrobacterium tumefaciens* induced SA accumulation. 24 h after infiltration, leaves were collected and metabolites were extracted as described for metabolite fingerprint analysis. Infiltration medium was used as mock treatment. The SA content was analyzed by LC-MS. Bars represent the mean ± STD of three biological replicates. Statistical differences among replicates are labeled with different letters ($P < 0.05$, one-way ANOVA and post hoc Tukey’s Test; $n = 3$).

| | LOCUS | NAME | FUNCTION | Supportability | MR |
|---|---|---|---|---|---|
| 0 | At5g67160 | EPS1 | HXXXD-type acyl-transferase family protein | 3 | 0 |
| 1 | At2g26170 | MAX1 | cytochrome P450, family 711, subfamily A, polypeptide 1 | 3 | 4.6 |
| 2 | At4g37430 | CYP91A2 | cytochrome P450, family 91, subfamily A, polypeptide 2 | 2 | 5.4 |
| 3 | At4g01070 | UGT72B1 | UDP-Glycosyltransferase superfamily protein | 3 | 5.6 |
| 4 | At4g33040 | ROXY21 | Thioredoxin superfamily protein | 3 | 6.6 |
| 5 | At2g44670 | DUF581 | Protein of unknown function (DUF581) | 2 | 9 |
| 6 | At4g29100 | DNA-binding | basic helix-loop-helix (bHLH) DNA-binding superfamily protein | 3 | 9.9 |
| 7 | At1g61930 | DUF584 | Protein of unknown function, DUF584 | 3 | 9.9 |
| 8 | At1g08250 | ADT6 | arogenate dehydratase 6 | 3 | 11.3 |
| 9 | At1g11700 | DUF584 | Protein of unknown function, DUF584 | 3 | 15 |
| 10 | At1g55210 | dirigent-like | Disease resistance-responsive (dirigent-like protein) family protein | 3 | 18.5 |
| 11 | At2g15310 | ARFB1A | ADP-ribosylation factor B1A | 3 | 19.2 |
| 12 | At3g20100 | CYP705A19 | cytochrome P450, family 705, subfamily A, polypeptide 19 | 3 | 21.5 |
| 13 | At4g36710 | HAM4 | GRAS family transcription factor | 3 | 21.9 |
| 14 | At2g21560 | nucleolar-like protein | nucleolar-like protein | 3 | 25.3 |
| 15 | At1g49740 | PLC-like phosphodiesterases | PLC-like phosphodiesterases superfamily protein | 3 | 26.9 |
| 16 | At1g06000 | UGT89C1 | UDP-Glycosyltransferase superfamily protein | 3 | 30.7 |
| 17 | At1g35910 | TPPD | Haloacid dehalogenase-like hydrolase (HAD) superfamily protein | 3 | 32.7 |
| 18 | At3g22550 | NAD(P)H-quinone oxidoreductase subunit, putative | NAD(P)H-quinone oxidoreductase subunit, putative | 3 | 34 |
| 19 | At3g14395 | MAT1 | Mite attack triggered immunity1 | 3 | 37.2 |
| 20 | At3g54390 | sequence-specific DNA binding transcription factors | sequence-specific DNA binding transcription factors | 3 | 37.9 |
| 21 | At3g15650 | alpha/beta-Hydrolases | alpha/beta-Hydrolases superfamily protein | 3 | 38.4 |
| 22 | At5g23580 | CPK12 | calmodulin-like domain protein kinase 9 | 2 | 40.2 |
| 23 | At3g21240 | 4CL2 | 4-coumarate:CoA ligase 2 | 3 | 42.9 |
| 24 | At1g28570 | SGNH hydrolase-type esterase superfamily protein | SGNH hydrolase-type esterase superfamily protein | 3 | 44 |
| 25 | At3g25290 | Auxin-responsive | Auxin-responsive family protein | 3 | 47.3 |
| 26 | At1g15950 | CCR1 | cinnamoyl coa reductase 1 | 3 | 47.7 |
| 27 | At1g32740 | SBP | SBP (S-ribonuclease binding protein) family protein | 2 | 49.7 |
| 28 | At4g34050 | CCoAOMT1 | S-adenosyl-L-methionine-dependent methyltransferases superfamily protein | 3 | 50.7 |
| 29 | At4g10390 | Protein kinase superfamily protein | Protein kinase superfamily protein | 3 | 54.8 |
| 30 | At4g30450 | glycine-rich protein | glycine-rich protein | 3 | 57.9 |
| 31 | At3g50630 | KRP2 | KIP-related protein 2 | 3 | 58.9 |
| 32 | At5g15910 | NAD(P)-binding Rossmann-fold superfamily protein | NAD(P)-binding Rossmann-fold superfamily protein | 3 | 59.3 |
| 33 | At5g63580 | FLS2 | flavonol synthase 2 | 3 | 59.5 |
| 34 | At5g44670 | GALS2 | Domain of unknown function (DUF23) | 3 | 66.1 |
| 35 | At4g36640 | Sec14p-like phosphatidylinositol transfer | Sec14p-like phosphatidylinositol transfer family protein | 3 | 66.7 |
| 36 | At5g37550 | hypothetical protein | hypothetical protein | 3 | 66.7 |
| 37 | At2g39830 | DAR2 | DA1-related protein 2 | 3 | 70 |
| 38 | At1g33260 | Protein kinase superfamily protein | Protein kinase superfamily protein | 3 | 70.9 |
| 39 | At4g22340 | CDS2 | cytidinediphosphate diacylglycerol synthase 2 | 2 | 72.9 |
| 41 | At2g41140 | CRK1 | CDPK-related kinase 1 | 3 | 73.7 |
| 40 | At3g52370 | FLA15 | FASCICLIN-like arabinogalactan protein 15 precursor | 3 | 73.7 |
| 42 | At5g46590 | NAC096 | NAC domain containing protein 96 | 3 | 74.1 |
| 43 | At3g16330 | | Avr9/Cf-9 rapidly elicited protein | 3 | 75.1 |
| 44 | At5g19875 | Transmembrane protein | Transmembrane protein | 3 | 78.1 |
| 45 | At5g17050 | UGT78D2 | UDP-glucosyl transferase 78D2 | 3 | 78.6 |
| 46 | At5g65280 | GCL1 | GCR2-like 1 | 3 | 79.1 |
| 47 | At1g72500 | heavy chain-like protein | heavy chain-like protein | 3 | 79.8 |
| 48 | At4g17880 | MYC4 | Basic helix-loop-helix (bHLH) DNA-binding family protein | 3 | 80.5 |
| 49 | At1g67360 | REF | Rubber elongation factor protein (REF) | 3 | 81.8 |
| 50 | At3g27170 | CLC-B | chloride channel B | 3 | 82.8 |
| 51 | At3g51240 | TT6 | flavanone 3-hydroxylase | 3 | 83.1 |
| 52 | At3g12750 | ZIP1 | zinc transporter 1 precursor | 3 | 85.2 |
| 53 | At2g41000 | DnaJ-domain | Chaperone DnaJ-domain superfamily protein | 3 | 86.8 |
| 54 | At1g16310 | Cation efflux | Cation efflux family protein | 3 | 87.7 |
| 55 | At3g21750 | UGT71B1 | UDP-glucosyl transferase 71B1 | 3 | 87.8 |
| 56 | At3g24170 | GR1 | glutathione-disulfide reductase | 3 | 90.2 |
| 57 | At4g34630 | prostatic spermine-binding-like protein | prostatic spermine-binding-like protein | 3 | 90.5 |
| 58 | At5g23760 | Copper transport | Copper transport protein family | 2 | 94.8 |
| 59 | At2g38180 | SGNH hydrolase-type esterase superfamily protein | SGNH hydrolase-type esterase superfamily protein | 2 | 96 |
| 60 | At3g13110 | SERAT2;2 | serine acetyltransferase 2;2 | 3 | 96.1 |
| 61 | At2g44300 | Bifunctional inhibitor/lipid-transfer protein/seed storage 2S albumin superfamily protein | Bifunctional inhibitor/lipid-transfer protein/seed storage 2S albumin superfamily protein | 3 | 99.5 |
| 62 | At5g02230 | HAD | Haloacid dehalogenase-like hydrolase (HAD) superfamily protein | 3 | 99.7 |
| 63 | At2g38710 | AMMECR1 | AMMECR1 family | 3 | 100.1 |
| 64 | At1g07640 | URP3 | Dof-type zinc finger DNA-binding family protein | 3 | 101.3 |
| 65 | At1g04220 | KCS2 | 3-ketoacyl-CoA synthase 2 | 3 | 103.5 |
| 66 | At1g28680 | HXXXD-type acyl-transferase family protein | HXXXD-type acyl-transferase family protein | 3 | 104.6 |
| 67 | At2g42760 | DUF1685 family protein | DUF1685 family protein | 3 | 104.8 |
| 68 | At2g46420 | helicase with zinc finger protein | helicase with zinc finger protein | 3 | 105.4 |
| 69 | At5g02270 | NAP9 | non-intrinsic ABC protein 9 | 3 | 105.8 |
| 70 | At4g39190 | nucleolar-like protein | nucleolar-like protein | 3 | 107.1 |

**Fig. S12.** Expression of *ICS1* (At1g74710), *EDS5* (At4g39030) and *PBS3* (At5g13320) is not co-regulated with *EPS1* (At5g67160). Co-expression analysis was performed using *EPS1* as bait with ATTED-II (http://atted.jp/; Version: 9.2). The Top 70 ranked co-expressed genes from microarray-based datasets are displayed. Immunity related genes are marked in yellow. Supportability is displayed by numbers (0-3) according to the p-value threshold (0=>1E-01, 1 =<1E-01, 2=<1E-01, 3=<1E-03) as a measure of reproducibility. The mutual rank (MR) index is indicated as a measure of co-expression (*9*).

### Table S1. Fragmentation data for investigated metabolites.

| | Type of ion | Elemental composition | Exact mass [Da] | MS/MS fragmentation | CE [eV] |
|---|---|---|---|---|---|
| **CA-7-Glu** | [M-H]- | C15H17NO9 | 355.090 | 336.072, 222.077, 163.04 | 8 |
| **CA-9-Glu** | [M-H]- | C15H17NO9 | 355.090 | 336.072, 216.051, 172.061 | 8 |
| **ISC-7-Glu** | [M-H]- | C15H17NO9 | 355.090 | 292.082. 274.072, 135.045 | 10 |
| **ISC-9-Glu** | [M-H]- | C15H17NO9 | 355.090 | 292.082, 216.051, 172.061, 137.024 | 10 |
| **SA** | [M-H]- | C7H6O3 | 138.032 | 93.034 | 10 |
| **2HNG** | [M-H]- | C8H11NO6 | 217.059 | 172.061, 136.040, 86.024 | 10 |
| **CA** | [M-H]- | C10H10O6 | 226.048 | 137.026, 121.029, 93.034 | 10 |
| **ISC** | [M-H]- | C10H10O6 | 226.048 | 137.026, 135.045, 119.050, 93.034 | 10 |

### Table S2. Primers used in this study.

| Primer | 5'-3' sequence | Purpose |
|---|---|---|
| ICS1-D47_EcoRI_fwd | ACGGAATTCATGAATGGTTGTGATGGAGA | cloning |
| ICS1-D47_HindIII_rev | ACGAAGCTTTCAATTAATCGCCTGTAGAGA | cloning |
| ICS1_w_Tag_SalI_fwd | ACGGTCGACATGGCTTCACTTCAATTTTCTTCT | cloning |
| ICS1-TAG-SalI_rev | ACGGTCGACGAAATCTCCATCACAACCATT | cloning |
| ICS1_TOPO_fwd | CACCATGGCTTCACTTCAATTTTCTTC | cloning |
| ICS1_TOPO_rev | ATTAATCGCCTGTAGAGATGTTG | cloning |
| PBS3_BamHI_fwd | ACGGGATCCATGAAGCCAATCTTCGATATCAACG | cloning |
| PBS3_HindIII_rev | ACGAAGCTTAATACTGAAGAATTTGGCTACCACAC | cloning |
| PBS3_TOPO_fwd | CACCATGAAGCCAATCTTCGATATC | cloning |
| PBS3_TOPO_rev | AATACTGAAGAATTTGGCTACCAC | cloning |
